# Profiling of intestinal T cells reveals antigen-specific dependency on early-life programming for the generation of microbiota-reactive Tregs

**DOI:** 10.64898/2026.09.23.753882

**Authors:** Lara Vetter, Jessica N. Witchley, So Hyun Ahn, R. Camille Brewer, Andrew Ly, Ali Tout, Allon Wagner, Gregory M. Barton

## Abstract

The generation of appropriate responses to the intestinal microbiota is an essential aspect of immune homeostasis. Studies of microbes that elicit stereotyped, antigen-specific CD4 T cell responses support the view that immune recognition of the microbiota is specific, relatively limited, and largely skewed toward Tregs, but recent profiling efforts have suggested that recognition of the microbiota may be broader and less regulatory. Such studies have mostly tracked responses to single antigens or used mice that acquire their microbiota after weaning, which has made it difficult to generalize the rules that govern microbiota-mediated T cell responses. Here we comprehensively profile, for the first time, the T cell response to the microbiota at homeostasis, and develop a panel of tools to track multiple, endogenous microbiota-reactive CD4 T cell populations in vivo. Our profiling reveals a broad microbiota-reactive T cell response that is Treg-biased but that also contains non-Treg responses to specific microbes. We find that certain microbiota-reactive T cells require priming during early-life to efficiently establish their Treg identity, while others do not. Our findings illustrate an unappreciated antigen-specific complexity in programming of microbiota-reactive T cells and underscore the importance of studying responses to the microbiota under homeostatic conditions.

## INTRODUCTION

The immune system has evolved to detect and eliminate pathogens, yet at densely colonized mucosal surfaces it must also tolerate innocuous microbial species. How this tolerance is established and how its breakdown gives rise to pathogenic immune responses remains a central question in mucosal immunology and is relevant to a range of chronic inflammatory diseases^1–3^. Over the last two decades, efforts to understand how the microbiota shapes host immunity have identified the induction of CD4+ T cells that recognize the microbiota as central contributors to tolerance. However, the rules governing how this tolerance is generated remain incompletely understood, and several basic questions persist: how broad is the recognition of the microbiota during homeostasis, is the response to a bacterium specific, and do the resulting phenotypes persist or shift when tolerance is perturbed?

To date, efforts to understand how the microbiota shapes host immunity have identified only a handful of bacterial species that elicit antigen-specific CD4+ T cell responses^4–12^, leading to the idea that most members of the microbiota may not be recognized, and that if they are, the response is narrowly directed against antigens unique to that species. However, recent work has suggested an alternative paradigm, in which recognition of the microbiota is broader than previously appreciated and mediated largely through epitopes conserved across taxa, rather than through the strain-unique antigens described above^13^. Thus, whether recognition of the microbiota concentrates on a handful of dominant organisms or is distributed broadly across most or all its members remains unclear^2^.

A substantial body of work has demonstrated that resident gut bacteria contribute to intestinal tolerance through the induction of regulatory T cells (Tregs). Antigen-specific recognition of individual bacterial species is emerging as an important route to Treg induction, although Tregs can also develop through microbial signals without direct TCR engagement. Tolerogenic Treg induction at steady state has been tied to recognition of defined Clostridia taxa^5,14,15^, Bacteroidetes species^16,17^ and Helicobacter hepaticus ^8,18^, while other bacterial species have been shown to elicit antigen-specific CD4+ T cell responses that adopt non-Treg fates instead^4,7,9,10,12,19–21^. Departures from a Treg-dominant outcome are typically explained as a property of a specific pathobiont^10^ or as a consequence of an already-inflamed environment, in which the antigen-specific repertoire constricts around a narrower, more pro-inflammatory set of specificities^22,23^.

Whether these homeostatic antigen-specific responses remain stable when tolerance is challenged is similarly unresolved. Acute inflammatory perturbations, whether infectious or chemically induced, have each been associated with a shift away from regulatory phenotypes toward more inflammatory effector fates, both at the population level^24,25^ and, in select cases, at the level of individual antigen specificities^6,18,23^. Yet these studies differ widely in the type of challenge used, the developmental stage of the host, and whether bulk or antigen-specific populations were tracked. Colonization occurring outside the early-life window represents a further, distinct challenge to tolerance because many tolerogenic responses depend on being exposed to the microbiota during the early life window^26–30^. Whether instability under perturbation is a general property of microbiota-specific T cell responses, or instead depends on the nature of the challenge, the maturity of the response, or the individual antigen involved, remains unclear, and resolving it requires tracking the same antigen-specific populations across multiple, distinct perturbations directly, something no study has yet done across a broad, natural microbiota.

These diverse findings on how microbiota-reactive T cells behave may reflect real biological differences rather than apparent contradictions, yet progress on the rules governing this response has been limited by technical challenges. Recent efforts to bridge this gap have leveraged high-throughput TCR synthesis and screening platforms capable of constructing and functionally testing many TCRs in parallel^13,13,31–33^; however, identifying naturally occurring gut T cells reactive to the microbiota remains a challenge. This difficulty stems in part from the sheer diversity of both the host TCR repertoire and the bacterial taxa that colonize the gut^34,35^, many of which are not culturable, making it nearly impossible to comprehensively test all candidate antigens against all mature T cells. Previous work aimed at circumventing these challenges has either used mice with a restricted TCR repertoire^36^, which narrows the antigens that can be recognized, or mice with reduced antigen diversity^37,38^, which fails to capture crosstalk between microbes. More recent work has characterized responses to more complex communities, but these studies have relied on human-derived rather than mouse-native strains and on horizontal colonization of adult mice^13,39,40^, which bypasses the early life weaning window in which critical host-microbe immune interactions are taking place and peripheral tissues are being seeded by peripherally induced Tregs required for tolerance^26–28^. In summary, each of these approaches has limitations, and no study to date has captured the nature of the homeostatic, endogenous response made against the microbiota in a comprehensive way.

Here, we took advantage of a fully defined, culturable 12-species microbiota (Oligo-Mouse-Microbiota, OMM12)^41^ to screen antigen-specific reactivity against every community member individually in vertically colonized mice. We used mice with an unrestricted TCR repertoire, combining single-cell RNA and TCR sequencing of endogenous CD4+ T cells with a scalable platform for reconstituting single-cell-derived TCRs as functional reporter lines. This approach revealed that recognition of the microbiota at homeostasis is broad and dominated by strain-unique rather than conserved antigens, that each species imprints a distinct T helper phenotype on its cognate T cells, and that the response is nonetheless overwhelmingly Treg-biased when considered across the whole community. Using antigen-specific tetramers generated against a subset of these epitopes, we found that pre-established, homeostatic Treg responses are largely resistant to acute inflammation, but that colonization timing can uncouple antigen recognition from Treg commitment in an antigen-specific manner. Together, these results indicate that the rules governing tolerance to the microbiota act at the level of the individual antigen rather than uniformly across the population, and that resolving them requires tracking many such responses in parallel across an intact community.

## RESULTS

### Single-cell transcriptomic and TCR profiling reveals broad, Treg-biased recognition of the microbiota at homeostasis

To characterize the T cell response against a microbiota under conditions in which tolerance is established under normal, physiological conditions, we maintained gnotobiotic C57BL/6N mice vertically colonized with the OMM12 community, rather than introducing the bacteria in adulthood, and analyzed the CD4+ compartment at homeostasis. The OMM12 microbiota is a fully defined, culturable 12-species bacterial consortium spanning five phyla (Verrucomicrobia, Bacteroidetes, Firmicutes, Proteobacteria, and Actinobacteria) that is representative of a normal SPF microbiota^41^.Throughout this study, we refer to the individual OMM12 strains by their standard abbreviations (e.g., Amuc, Bcae, Mint, Cinn, Lreu, Efae, Eclo, Bpse, Fpla, Amur, Tmur, Bani). Importantly, the OMM12 community stably colonizes the murine intestine over many generations, allowing every member to be reliably transmitted from mother to pup and to be present during the early life window^42^.

We isolated antigen-experienced (CD44+CD4+TCRβ+) T cells by fluorescence-activated cell sorting (FACS) from the gut-associated lymphoid tissues (mesenteric lymph nodes (MLN) and Peyer’s patches (PP)) and lamina propria (small and large intestine (SILP and LILP)), and generated paired single-cell RNA and TCR sequencing (scRNA-seq and scTCR-seq) libraries from these populations (Figure 1A). Unsupervised clustering of the full dataset resolved 17 transcriptionally distinct populations, annotated using canonical lineage-defining transcripts (Figure S1A–C), with every cluster represented in all mice. Because our goal was to relate microbiota-reactive TCRs to broad functional lineages rather than finely subdivided transcriptional states, we collapsed these clusters into nine broad T helper (Th) phenotypes (Figure 1B) that were used throughout the TCR-specificity analyses.

**Figure 1.**
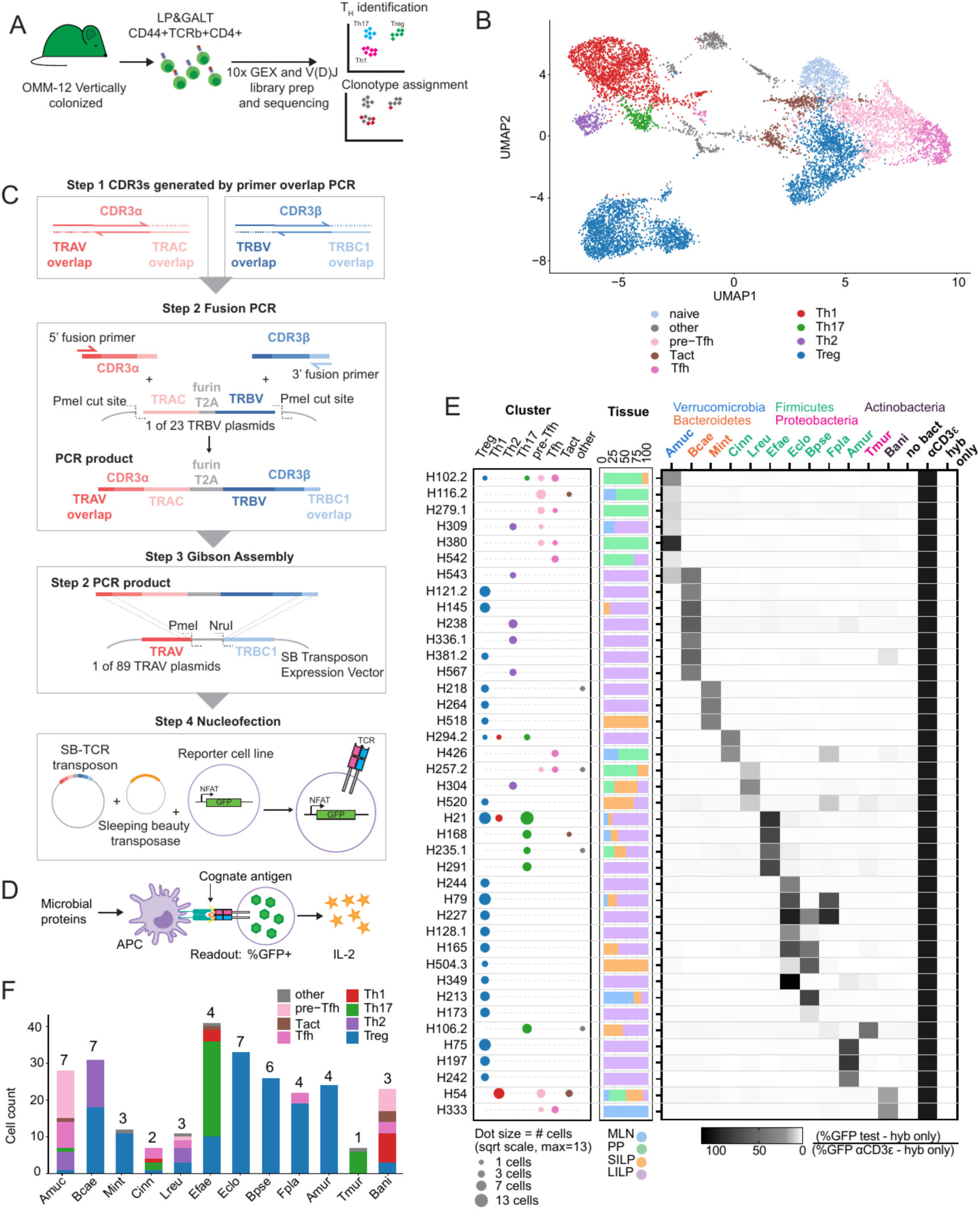
scRNA-seq and scTCR-seq reveal Treg-biased microbiota recognition at homeostasis. (A) Schematic of workflow to generate scRNA-Seq and scTCR-seq datasets. CD44+CD4+TCRb+ T cells were sorted from the gut-associated lymphoid tissues (GALT) and lamina propria (LP) of vertically-colonized OMM12 mice (N=2 males) and used to generate 10x scRNA-seq and scTCR-seq libraries. (B) UMAP of the broad CD4 helper T cell clusters identified by scRNA-seq in (A) and applied to reactivity data shown in (E) and (F). N=10,020 cells. (C) Schematic of workflow to assemble and express TCR genes. Sleeping beauty transposon plasmids containing single-cell derived TCRs were made using a combination of (Step 1) overlap PCR, (Step 2) fusion PCR and (Step 3) Gibson assembly. TCRs were stably integrated into an NFAT-GFP reporter cell line using nucleofection(Step 4). (D) Reactivity screening of TCR-expressing reporter cells. Cells were cocultured with CD11c+ antigen-presenting cells (APC) and different antigen sources (individual heat-killed OMM12 microbes, germ-free or OMM12 colonized lumenal cecal contents, or aCD3 antibody) for 12-24 h. TCR activation was determined by flow cytometry (% GFP+) or IL-2 ELISA. (E) Compilation of Th phenotype (left), tissue origin (middle) and reactivity (right) data for T cells reactive to individual members of the OMM12 microbiota. TCR clonotype numbers are indicated on the left axis.(F) Summary of data from (E). The number of TCRs identified with reactivity for each OMM12 member is indicated above each bar. MLN = mesenteric lymph node, PP = Peyer’s patch, SILP = small intestine lamina propria, LILP = large intestine lamina propria; OMMCe = cecal contents from OMM12 colonized mice, GFCe = cecal contents from germ-free mice.

To link antigen specificity with transcriptional phenotype at scale, we developed a pipeline to rapidly assemble, express, and screen the reactivity of TCRs identified from our scRNA-seq dataset (Fig. 1C). Briefly, paired TCRα/β sequences were reconstituted into Sleeping Beauty transposon plasmids using a modular cloning strategy adapted from a human TCR cloning system^43^. We constructed 89 plasmids containing individual murine TRAV genes and 23 plasmids containing individual murine TRBV genes, each linked to TRBC1 or TRAC genes, respectively. CDR3 sequences were generated by overlap extension PCR (Figure 1B, Step 1) and then combined with the relevant TRAV and TRBC plasmids via Fusion PCR and Gibson assembly (Figure 1B, Steps 2 and 3). Verified TCR-expressing plasmids were introduced by nucleofection into a CD4⁺/CD3⁺ 58α⁻β⁻ hybridoma line engineered for stable expression of murine CD4, CD3, and an NFAT-GFP reporter (Figure 1C, Step 4), generating 292 TCR-expressing hybridomas that were then tested in an activation assay in which dendritic cells (DCs) and hybridomas were mixed with each individual OMM12 strains, as well as germ-free (GF) and OMM12 cecal contents and stimulated overnight (Figure 1D). Of all clonotypes screened, ^73^ showed reactivity (∼25%), with 41 responding to at least one OMM12 bacterial species (Figure 1E). Surprisingly, we identified microbe-reactive clonotypes against every member of the OMM12 consortium, spanning all five constituent phyla (Figure 1E and 1F, Figure S1D); the largest number of reactive clonotypes mapped to Firmicutes species. The great majority of reactive TCRs recognized only a single species, indicating that recognition is dominated by private, non-cross-reactive specificities. 11 clonotypes reacted to multiple species, five clonotypes were multi-strain reactive within Firmicutes, and one clonotype (H543) was cross-reactive across phyla, recognizing both Amuc and Bcae (Figure 1D).

To determine whether TCR reactivity to a given bacterial species was associated with a particular Th phenotype, we classified reactive clones by their phenotypic profile (Figure 1F). Reactivity to individual species was associated with a largely species-specific distribution of T helper phenotypes. Clones reactive to Eclo, Bpse, Mint, Amur and Fpla-reactive clones were entirely or almost entirely Treg, while Bcae-reactive clones split between Treg and Th2 phenotypes. In contrast, Amuc-reactive clones were enriched for Tfh and pre-Tfh phenotypes, consistent with our previous work characterizing the response to this bacterium7, and Efae-and Tmur-reactive clones were dominated by Th17. The remaining species (Cinn, Lreu, and Bani) yielded more heterogeneous profiles spanning multiple phenotypes (Treg, Th2, Th17, Th1, Tfh, and pre-Tfh). We also confirmed that Il10 was not enriched within our Th1 clusters, distinguishing this population from a Tr1-like phenotype recently described in the small intestine (Figure S1A)^44^. Strikingly, the vast majority of species (11/12) skewed toward Treg induction, revealing an unexpectedly strong Treg bias in the response.

We identified 11 additional clonotypes that reacted to OMM12 cecal contents, but for which we could not identify reactivity with any individual OMM12 members (Figure S2A). These TCRs likely recognize antigens that are not expressed under the culture conditions used to propagate OMM12 members in vitro. Five hybridomas reacted to all conditions tested, including the no bacteria control, suggesting these are likely self-reactive TCRs that recognize antigens expressed by the DCs used in the screening assay (Figure S2A).

Additionally, our screening identified TCRs that were non-microbiota reactive. We identified 16 TCRs that responded to both OMM12 and GF cecal contents, suggesting reactivity to chow (Figure S2). We rescreened these TCRs against components of the chow and previously identified dietary antigens^45–48^. Several of these TCRs recognized dietary antigens present in standard chow, including maize zein, soy glycinin, and wheat gliadin peptides, indicating that our screen also captures food-antigen-reactive clones (Figure S2)^47,48^.

Overall, our findings from this screen uncovered that during homeostasis the majority of the response to the microbiota is tolerogenic, although individual bacteria can stereotype the response that is elicited by the immune system.

### Acute intestinal inflammation broadens Th17 reactivity across the bulk repertoire while leaving established antigen-specific phenotypes largely intact

Having found that most microbiota-specific T cells adopt a Treg phenotype under homeostatic conditions, we wanted to know whether this broad tolerogenic bias is maintained under inflammatory conditions. Tregs are required to keep the immune system from reacting against innocuous microbes, and their loss of function is thought to contribute to IBD^47,49,50^; however, most studies of microbiota-reactive T cells during inflammation have relied on transferring naïve cells into colitis-prone hosts^51–53^, leaving open the question about how a response already shaped by homeostasis is affected once inflammation is induced. We therefore asked how acute intestinal inflammation reshapes the antigen-specific landscape we had just characterized.

Vertically colonized OMM12 mice were fed chow supplemented with piroxicam (px), a nonsteroidal anti-inflammatory drug (NSAID) that transiently disrupts the epithelial barrier, for 10 days together with disruption of the IL-10 circuit via administration of an anti-IL-10 receptor (αIL-10R) blocking antibody (Figure 2A)^25,54–57^. This combined insult produced an inflammatory episode in the intestine: fecal lipocalin-2 (Lcn-2)^58^ rose in mice treated with px+αIL-10R and remained elevated throughout the regimen (Figure 2B). Profiling of the small intestinal lamina propria at day 17 showed an expanded Th17 population (Figure 1C) as well as an expansion of the myeloid compartment in the LP (Figure S3A). Mice treated with px+αIL-10R also exhibited increased spleen weight at day 17 (Figure 1D). In contrast, the Treg compartment remained unchanged by the end of the treatment, indicating that this combined insult selectively expands Th17 cells without depleting the pre-existing Treg compartment (Figure S3A).

**Figure 2.**
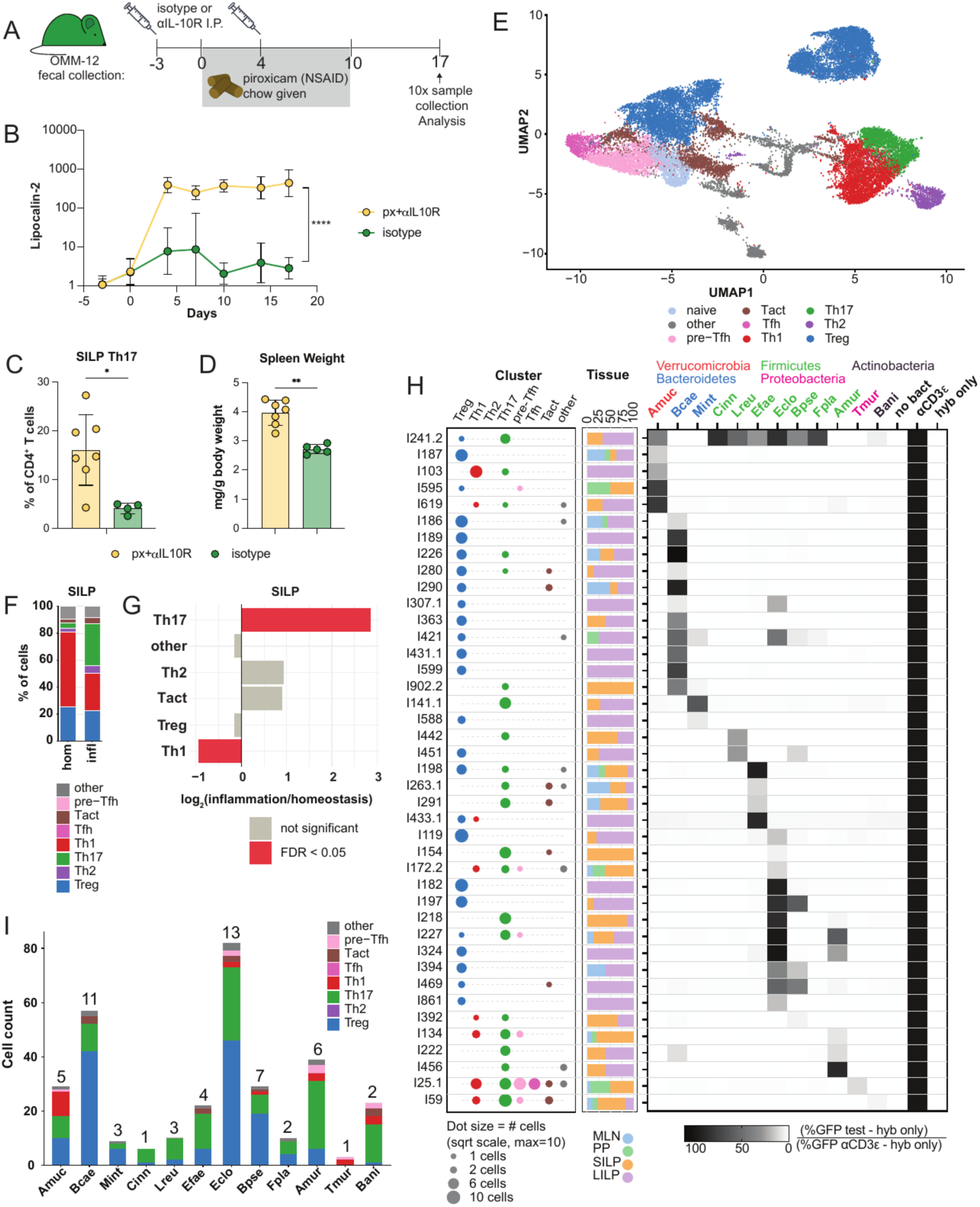
Microbiota recognition after acute inflammation. (A) Schematic of the experimental timeline used to induce acute intestinal inflammation. Vertically colonized OMM12 mice were given piroxicam-supplemented (px) chow ad libitum for 10 days in combination with 1 mg ɑIL-10R antibody or isotype control antibody by intra-peritoneal(IP) injection at days −3 and 4. Feces were collected at days -3, 0, 4, 7, 10 and 17 relative to being given the px chow. (B) Fecal Lipocalin-2 as measured by ELISA from the indicated groups of mice treated as described in (A). Geometric mean ± geometric SD are shown. (N=7 px+ɑIL-10R, N=5 isotype). P values determined by Mixed effect model with Greenhouse-Geisser Correction and Dunnett’s multiple comparison test with variance computed for individual comparisons at each time point separately. ****p < .0001. (C) Frequency of Th17 cells from the small intestine lamina propria (SILP) for mice in treatment groups from (B) at d17. Mean ± SD are shown. P values determined by two-tailed Mann-Whitney test, *p < .05. (D) Spleen weights for mice in treatment groups from (B) at d17. Mean ± SD are shown. P values determined by two-tailed Mann-Whitney test, **p < .01. (E) UMAP highlighting the broad CD4 Th phenotypes identified by analysis of the integrated single cell sequencing datasets from homeostasis and inflammation (N=24,117 total cells; 10,020 from homeostasis and 14,097 from inflammation). CD44+CD4+TCRβ+ T cells were sorted from the GALT and LP of vertically-colonized OMM12 mice (N=2) at d17 and used to generate 10x scRNA-seq and scTCR-seq libraries. (F) Proportion of cells in each cluster shown in (E) from the SILP at homeostasis (N=2,722) and inflammation (N=3,767). (G) Fold enrichment and false discovery rate (FDR) from propeller analysis of the broad clusters for the SILP. (H) Compilation of Th phenotype (left), tissue origin (middle) and reactivity (right) data for T cells reactive to individual members of the OMM12 microbiota. (I) Summary of data from (H). The number of TCRs identified with reactivity for each OMM12 member is indicated above each bar.

To capture how this acute inflammatory episode reshapes the antigen-specific T cell compartment, we repeated our scRNA/TCR-seq on GALT and LP antigen-experienced CD4+ T cells at day 17 and integrated the data with the homeostasis dataset (Figure 2E), which we again collapsed into the same T helper phenotypes based on identity of the clusters (Figure S3B-C). Cells from both the homeostasis and inflammation conditions populated every one of these clusters, indicating that inflammation does not generate qualitatively new CD4+ T cell states but instead redistributes cells among the phenotypes already present at homeostasis. Compositional analysis of the SILP showed a significant increase in the Th17 fraction and a significant decrease in the Th1 fraction after px+αIL-10R treatment (Figure 2F and 2G). This increased Th17 frequency is consistent with our flow cytometry data (Figure 2C).

Next, we used our established TCR assembly and screening platform to test the reactivity of 186 validated TCRs from this inflammation dataset. In addition to screening the top 50 most abundant TCR clonotypes, we prioritized testing the reactivity of LP pTregs, which had shown the greatest and broadest microbiota reactivity among the homeostasis dataset, and of Th17 cells, which expanded substantially during inflammation. This screening resulted in a ∼39% overall reactivity rate, which is notably higher than the ∼25% overall reactivity rate observed for the entire homeostasis T cell dataset; however, this increase is more likely a reflection of our selective screening of Tregs and Th17 cells. Indeed, when considering Tregs and Th17 cells in isolation, the reactivity rates for both cell types were essentially unchanged between the inflamed and homeostasis datasets (LP pTreg: 48.1% at homeostasis vs. 44.9% at inflammation; Th17: 23.4% at homeostasis vs. 20.0% at inflammation). Independent of screening strategy, Th17 clonotypes also made up a substantially larger share of the most abundant clonotypes overall during inflammation, whether measured by clonotype (38.0% vs. 82.0% of the top 50) or by cell number (11.2% vs. 38.7% of cells within the top 50). These numbers confirm that the Th17 compartment genuinely expands with inflammation, yet the frequency of microbiota-reactive Th17 cells remains the same between homeostasis and inflammation.

Of the TCRs that showed reactivity, 41 reacted to at least one OMM12 bacterial species; 31 reacted to a single species, and 10 reacted to more than one species. We again observed reactivity to all 12 members of the OMM12 consortium. Within this targeted screen, Treg-phenotype reactivity remained detectable in the majority of OMM12 species (11 of 12), indicating that Treg reactivity to the microbiota persists broadly even under our Th17/pTreg-enriched screening approach. We identified 19 additional TCRs that reacted to both OMM12 and GF cecal contents, and 10 TCRs that reacted to OMM12 cecal contents alone (Figure S4A). Screening the TCRs in the former category against chow components confirmed that they are food-reactive (Figure S4B). Notably, T cells reactive to αZein, which were exclusively Tregs at homeostasis, now also included a small fraction of effector-phenotype cells (Figure S4). Two clones reacted to all conditions tested, again likely indicating self-reactivity (Figure S4A).

Overall, these findings demonstrate that microbiota reactivity remains broad during inflammation, and that Treg recognition of the microbiota persists across most OMM12 species. We observe expansion of the Th17 compartment with inflammation, as well as Th17 reactivity with more OMM12 species. It is difficult to draw definitive conclusions about any change in Th17 reactivity from these data, though, because of the static nature of the information gained from such profiling pipelines. Our screening data cannot resolve whether the microbiota-reactive Th17 clonotypes we recovered in our inflammation dataset represent the conversion of previously Treg-committed antigen specific clones, the expansion of pre-existing but minor Th17 clones, or the recruitment of new specificities primed for the first time during inflammation. To make these distinctions requires tracking of cells with defined antigen specificities across different contexts. Therefore, we next sought to identify the specific antigens recognized by microbiota-reactive T cells from our profiling pipeline and to generate reagents to track such cells directly.

### Genomic library screening identifies mostly private and some shared antigens recognized by microbiota-reactive TCRs

To identify the antigens recognized by specific microbiota-reactive T cells and ultimately develop tools to track such cells in vivo, we selected five OMM12 members representative of the patterns of reactivity revealed by our profiling studies: Amur and Eclo (which were exclusively Treg at homeostasis but switched to a mixed Treg/Th17 phenotype at inflammation), Bcae (Treg biased in both screens), Efae (Th17 biased in both screens) and Bani (heterogeneous response). We screened genomic DNA expression libraries from these five OMM12 species^59,60^ against pools of TCR hybridomas identified as reactive against the relevant OMM12 member. Following the workflow outlined in Figure S5A, we identified 13 I-Ab binding peptides across the five OMM12 species that stimulate specific microbiota-reactive T cells (Table 1). Peptide titrations confirmed dose-dependent activation of the corresponding hybridomas for each peptide (Figure 3A-3G).

**Figure 3.**
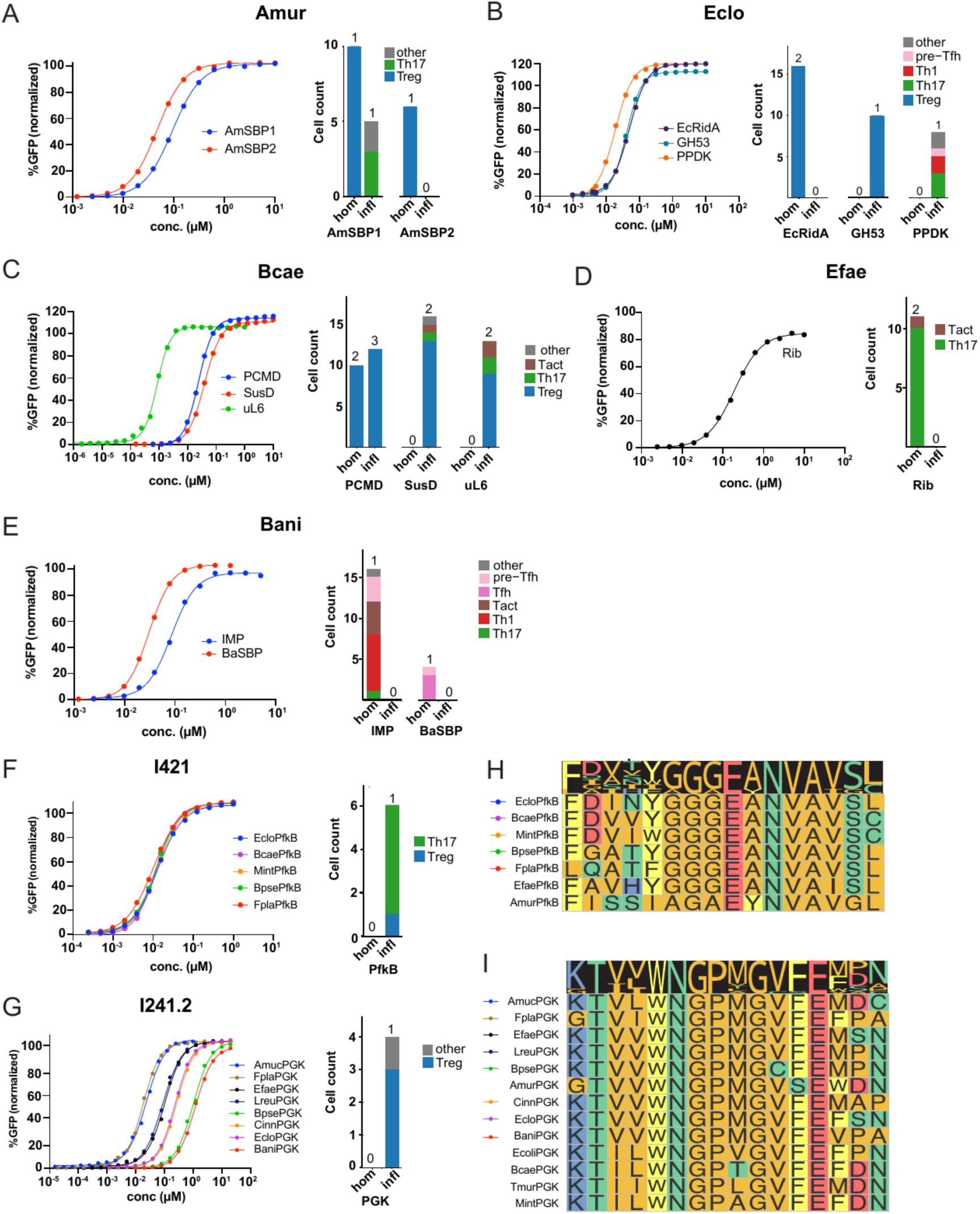
Identification of epitopes for microbiota-reactive TCRs in OMM12 mice. Genomic DNA libraries from Amur, Eclo, Bcae, Efae, and Bani were screened against TCRs reactive to each to identify cognate epitopes. Reactive TCR-expressing hybridomas were cocultured with mutuDC1s and cognate antigens. (A-G). Dose response curves (left) and summary of T cell phenotypes of peptide-reactive T cells from the combined homeostasis and inflammation single-cell profiling data (right) for Amur-(A), Eclo-(B), Bcae-(C), Efae-(D), and Bani-(E) reactive TCRs. Lines on dose response curves are sigmoidal four-parameter logistic (4PL) model fits of peptide concentration. The number of TCRs that contribute to the stacked bar plots is given above each bar. (F) Dose response curves (left) of I421 stimulated with peptides from all identified orthologs of PfkB from organisms that the TCR is reactive to as whole bugs. Those not shown were not stimulatory. Single cell annotated phenotypes(right) of cells that express the I421 TCR. (G) Dose response curves of I241.2 stimulated with peptides from all identified orthologs of PGK from organisms that stimulate the TCR as whole bugs. Those not shown were not stimulatory. Single cell annotated phenotypes(right) of cells that express the I241.2 TCR. (H) Alignment of all identified orthologs of PfkB from OMM12. Order of sequences indicates similarity to EcloPfkB. Some OMM12 members did not have an ortholog. (I) Alignment of all identified orthologs of PGK in OMM12 and Ecol. Order of sequences indicates similarity to AmucPGK.

**Table 1.**
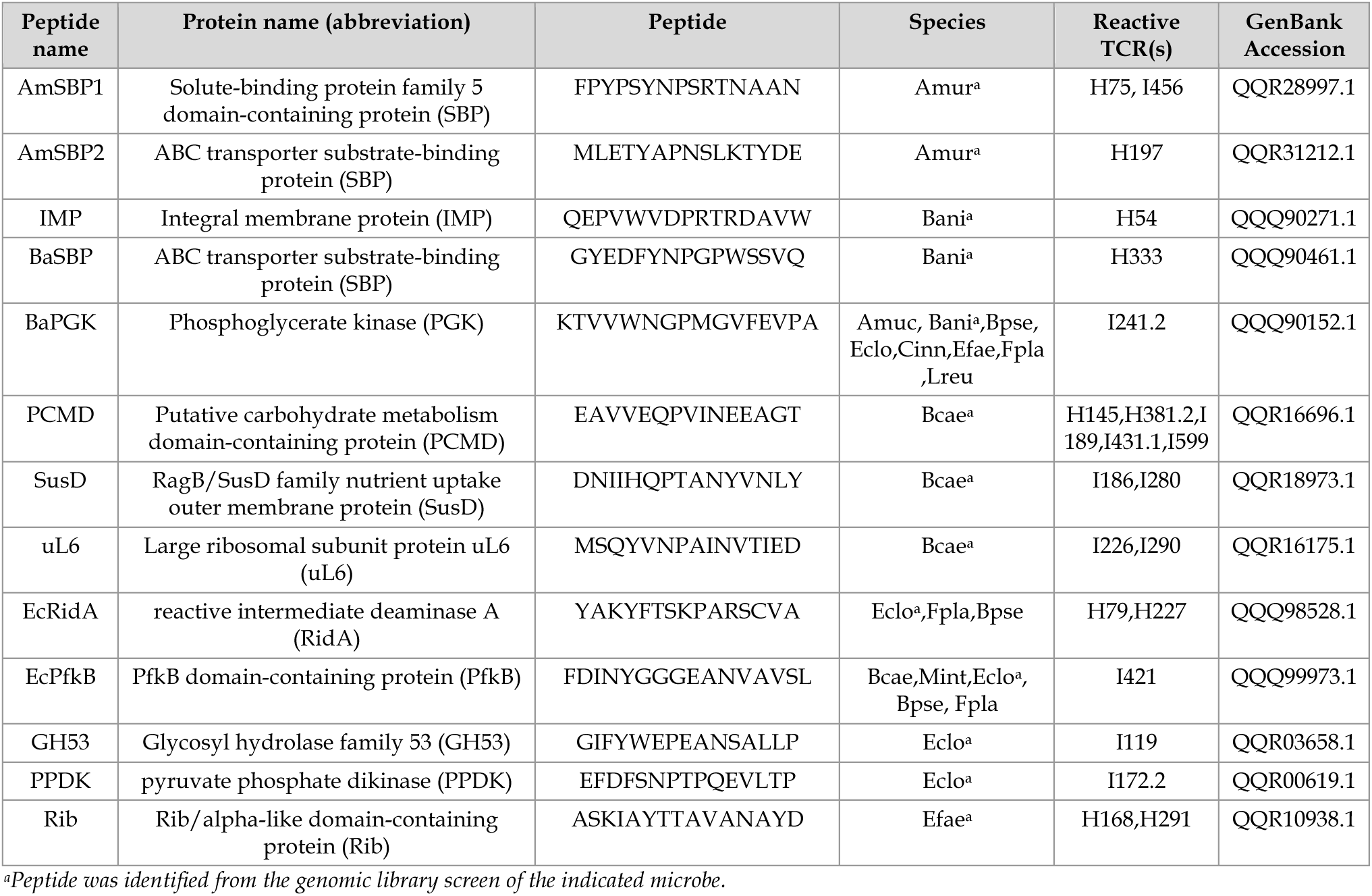
| Identified peptide sequences and additional details. Genomic DNA library screens were performed for Amur, Bani, Bcae, Eclo and Efae and the peptides that are validated to stimulate at least one TCR are listed.

The epitopes we identified were strikingly diverse and largely did not converge on shared antigens across species. Ten of the thirteen validated epitopes were strain-private, recognized only by TCRs reactive to their species of origin, and included both predicted cytosolic proteins (e.g., PPDK, uL6) and predicted surface-exposed or secreted antigens (solute-binding proteins (SBPs), outer membrane transporters, and cell envelope-anchored proteins). The remaining three antigens we identified, BaPGK, EcRidA, and EcPfkB, were cross reactive across multiple OMM12 species, and notably all three derived from conserved, cytosolic housekeeping proteins. Titration of the orthologous peptides from other OMM12 species confirmed that this cross-reactivity tracked with sequence conservation. Reactivity of the PfkB- and PGK-specific hybridomas (I421 and I241.2) against synthesized orthologs from across the OMM12 community correlated with the degree of sequence identity to the reference peptide, with the most divergent orthologs failing to stimulate the cognate hybridoma (Figure 3H-I). Together, these results suggest that cross reactivity in our dataset tracks how conserved a given peptide is across species, rather than reflecting a promiscuous TCR in these cases.

In general, the phenotype of the T cells reactive with specific peptides recapitulated the patterns we observed in our screening of whole bacteria. For example, T cells specific for Amur (AmSBP1, AmSBP2), Eclo (GH53, EcRidA), and Bcae (PCMD, SusD, uL6) were predominantly Treg at homeostasis (Figure 3A-3C, right panel), and T cells specific for Efae (Rib) were predominantly Th17 (Figure 3D, right panel). These phenotypes are consistent with the reactivity pattern we saw for these species by whole-bacteria screening during homeostasis. With inflammation, AmSBP1-specific T cells shifted from Treg to predominantly Th17, and the cross-reactive PfkB-specific T cells were also largely Th17, mirroring the shift toward Th17 reactivity we observed during inflammation (Figure 2I). Bcae- SusD- and uL6-specific T cells took on a mixed Treg/Th17 phenotype during inflammation that was still Treg-biased (Figure 3C, right panel), PPDK-specific T cells were split across Th17, Th1, and pre-Tfh (Figure 3B, right panel), while IMP- and BaSBP-specific T cells at homeostasis were predominantly Th1 and Tfh/pre-Tfh, respectively (Figure 3E, right panel), reflecting the same phenotypic heterogeneity present in the whole-bacteria reactive pool for Bani.

For most of the peptides, we only identified reactive TCRs from either the homeostasis or the inflammation dataset, which precluded us from drawing firm conclusions about how inflammation may shape the response; however, in two cases, PCMD and AmSBP1, TCRs specific for the same peptide were independently recovered from both datasets. This allowed us to compare cells specific for the same antigen before and after inflammation even though the two datasets were derived separately and from different animals. The PCMD-specific cells were all Tregs in both conditions (Figure 3C, right panel), whereas AmSBP1-specific cells shifted from predominantly Treg at homeostasis to predominantly Th17 during inflammation (Figure 3A, right panel). The differing effect on these two T cell populations makes it challenging to draw any general conclusions about how inflammation shapes the microbiota-reactive T cell response and underscores the limitations of scRNA-Seq-based profiling approaches. Therefore, we moved to generate MHC class II tetramers that would enable us to track microbiota-reactive T cells in larger numbers of animals and across more contexts.

### Established microbiota-specific T cell phenotype is largely stable despite acute intestinal inflammation

To enable direct tracking of OMM12-reactive T cells, we generated MHC class II tetramers against ten of the validated epitopes. We confirmed that each tetramer specifically stained its cognate hybridoma (Figure S6A–C). When we used these tetramers to stain endogenous CD4+ T cells from OMM12-colonized mice, six produced tetramer+ populations that were significantly enriched above background. The remaining tetramers did not label a population distinguishable from background staining, likely reflecting the low frequency and/or low avidity of the corresponding endogenous T cell responses in vivo. We therefore focused our subsequent analyses on the six tetramers that showed reproducible in vivo staining: PCMD, SusD, and uL6 from Bcae; GH53 and RidA from Eclo; and IMP from Bani. For Bcae, we initially characterized cells using tetramers against PCMD, SusD, and uL6 individually (Figure S6D-G); however, because all three tetramers stained populations with similar phenotypes, we pooled the Bcae tetramers for all subsequent experiments.

Initially we used Bcae, GH53, RidA, and IMP tetramers to track and characterize their corresponding T cell populations in mice at homeostasis. We analyzed various lymphoid tissues of GF, Altered Schaedler Flora (ASF)-colonized, and OMM12-colonized mice. For the EcRidA tetramer, we noted that its cognate peptide shares 93% sequence similarity with a peptide derived from ASF500, a member of the ASF community, so GF mice were used as the negative control for this tetramer instead. Bcae-, GH53-, and RidA-tetramer+ cells were readily detectable in the LILP of OMM12-colonized mice, and were essentially absent in GF and ASF controls, confirming their specificity for OMM12-derived antigens (Figure 4A–4C). The majority of these Tregs were RORγt positive and localized in the LILP (Figure S6J), consistent with our scRNAseq findings. In contrast, IMP-tetramer+ cells were present in higher numbers in the SILP rather than the LILP (Figure 4D) which is consistent with the localization and phenotype of IMP+ cells from the scRNAseq profiling. Tetramer+ cells for Bcae, GH53, and RidA were overwhelmingly Foxp3+ in the LILP at homeostasis (Figure 4E-F), while IMP-tetramer+ cells in the SILP showed a more heterogeneous, Th1/Th17-dominant composition with a smaller Treg fraction (Figure 4G). These phenotypes largely matched the scRNA-seq-based predictions (Figures 1F and 2F).

**Figure 4.**
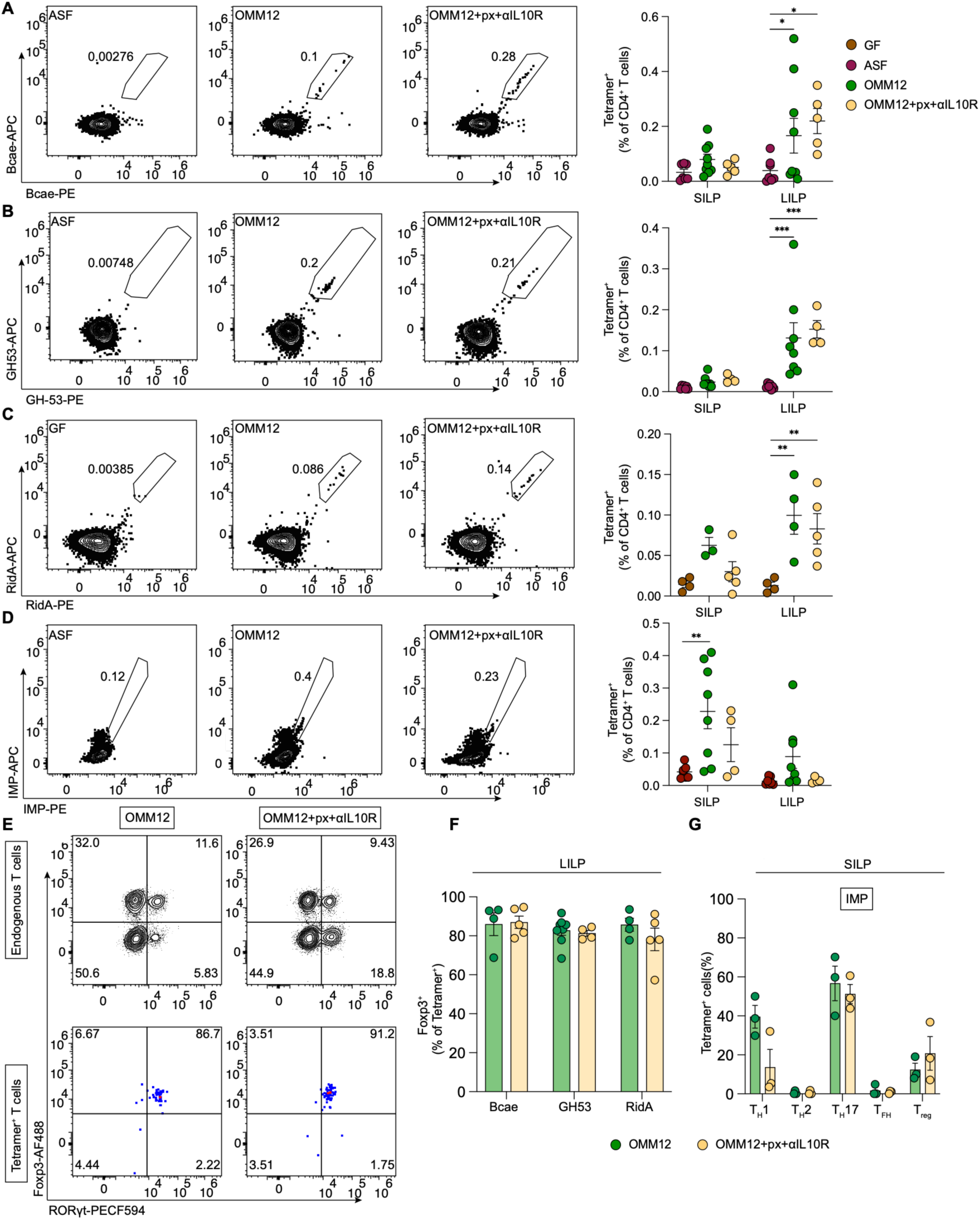
Piroxicam and IL-10R blockade minimally disrupts established microbiota-specific Treg identity. (A–D) Representative flow cytometry and summary plots of Bcae- (A), GH53- (B), RidA- (C), and IMP- (D) tetramer⁺ CD4⁺ T cells from the small intestinal (SILP) and large intestinal (LILP) lamina propria of germ-free (GF), Altered Schaedler Flora (ASF), OMM12-colonized, or OMM12-colonized mice treated with piroxicam plus anti-IL10R blocking antibody (OMM12+px+αIL10R). Numbers on representative plots indicate percent tetramer⁺ of CD4⁺ T cells. (E) Representative flow cytometry of Foxp3 and RORγt expression among tetramer⁻ endogenous (top) and Bcae-tetramer⁺ (bottom) CD4⁺ T cells from the LILP of OMM12 or OMM12+px+αIL10R mice. Quadrant values indicate the percentage of cells within each respective parent gate (endogenous or tetramer+ CD4+ T cells). (F) Summary of Foxp3⁺ frequency (% of tetramer⁺ cells) among Bcae-, GH53-, and RidA-specific CD4⁺ T cells in the LILP, comparing homeostasis (OMM12) to OMM12+px+αIL10R treatment. (G) T helper subset composition (Th1, Th2, Th17, Tfh, Treg; % of tetramer⁺ cells) among IMP-specific CD4⁺ T cells in the SILP. Bars show mean ± SEM; Each dot in summary plots represents an individual mouse; bars/lines indicate mean ± SEM. For (F) and (G), only mice with tetramer⁺ frequency exceeding the mean + 1 SD of the ASF control frequency were included (see Methods). p values calculated by two-way ANOVA with Sidak’s multiple comparisons test; *p < .05, **p < .01, ***p < .001.

Next we used our tetramer panel to examine whether T cells recognizing these epitopes changed during inflammation by subjecting mice to the px+αIL-10R regimen described earlier (Figure 2A). Tetramer+ populations for all four epitopes remained significantly enriched over GF/ASF background in mice that had undergone the intestinal inflammation regimen, but at frequencies not detectably different from untreated OMM12 mice (Figure 4A–4D), indicating that acute inflammation does not measurably alter the size of these antigen-specific populations. Phenotypically, Bcae-, GH53-, and RidA-tetramer+ cells in the LILP were overwhelmingly Foxp3+ in both homeostasis and after inflammation. This lack of change contrasted sharply to the bulk, tetramer-negative endogenous CD4+ T cells from the same mice, which showed an increase in Th17 frequencies (Figure 4E and 4F, Figure S6H-I). Similarly, the T helper subset composition of IMP-specific cells in the SILP, which consisted predominantly of Th1 and Th17 phenotypes with a smaller Treg fraction, was largely comparable between homeostasis and inflammation (Figure 4G), aside from a decrease in Th1 frequency, consistent with our earlier analysis of Th population changes during the px+αIL-10R regimen (Figure 2E and 2F). Overall, this analysis of microbiota-reactive Tregs using a panel of five MHC class II tetramers indicates that the phenotype of these cells is remarkably stable during acute inflammation with no evidence of a shift toward Th17 differentiation. These data suggest that the Th17 phenotype observed for some microbiota-reactive TCRs in the inflammation scRNA-Seq profiling dataset does not reflect a reprogramming of established microbiota-reactive Tregs that developed during homeostasis. Instead, it is possible that acute intestinal inflammation recruits new, largely non-overlapping TCRs specificities into the Th17 lineage.

### Horizontal colonization of adult germ-free mice partially uncouples antigen recognition from stable Treg commitment

Given the surprising stability of microbiota-specific Tregs during acute inflammation, we next considered whether any perturbation could alter their differentiation. Many microbiota-reactive Tregs are generated during an early-life window^26–30^, yet most studies of microbiota-reactive T cells, especially profiling studies performed at scale, are carried out in GF mice colonized as adults. These studies have typically reported a higher incidence of non-Tregs with microbiota reactivity^11,13,19,21,61^, so we reasoned that this difference may be due to colonization outside this critical window, and that having an already established Treg pool may inhibit non-Treg responses in an antigen specific manner.

To test this hypothesis directly, we tracked microbiota-reactive T cells using our panel of MHC class II tetramers in adult GF mice colonized horizontally with OMM12 (exGF>OMM12). Intestinal T cell responses were analyzed two weeks after colonization to be consistent with prior studies^13,61^ (Figure 5A). Bcae-, GH53-, RidA-, and IMP-specific populations were all detectable in exGF>OMM12 mice at frequencies comparable to, or in the case of GH53, exceeding, those seen in vertically colonized mice (Figure 5B-E). Strikingly, horizontal colonization altered the phenotype of these cells in an antigen-specific manner. Bcae- and RidA-tetramer+ cells in the LILP of exGF>OMM12 mice showed a significant reduction in Foxp3+ frequency and a reciprocal increase in the Th17 fraction relative to vertically colonized OMM12 mice (Figure 5F–5H). IMP-tetramer+ cells in the SILP also showed a significant increase in the frequency of Th17 cells (Figure 5F, 5I-J). In contrast, GH53-tetramer+ cells remained almost uniformly Foxp3+ regardless of whether OMM12 was acquired vertically or horizontally (Figure 5G and 5H). This antigen-specific shift in phenotype occurred against a backdrop of increased Th17 frequencies in the bulk CD4 T cell compartment in the SILP and LILP of exGF>OMM12 mice (Figure S9A). These findings suggest that horizontal colonization of adult mice with an intact, defined microbiota is sufficient to generate and expand microbiota-specific T cells but is not sufficient to reliably commit them to a fully differentiated Treg fate. Some antigens (Bcae, RidA) show a strong requirement for early-life exposure to establish durable Treg identity, while at least one (GH53) generates a Treg response irrespective of colonization timing.

**Figure 5.**
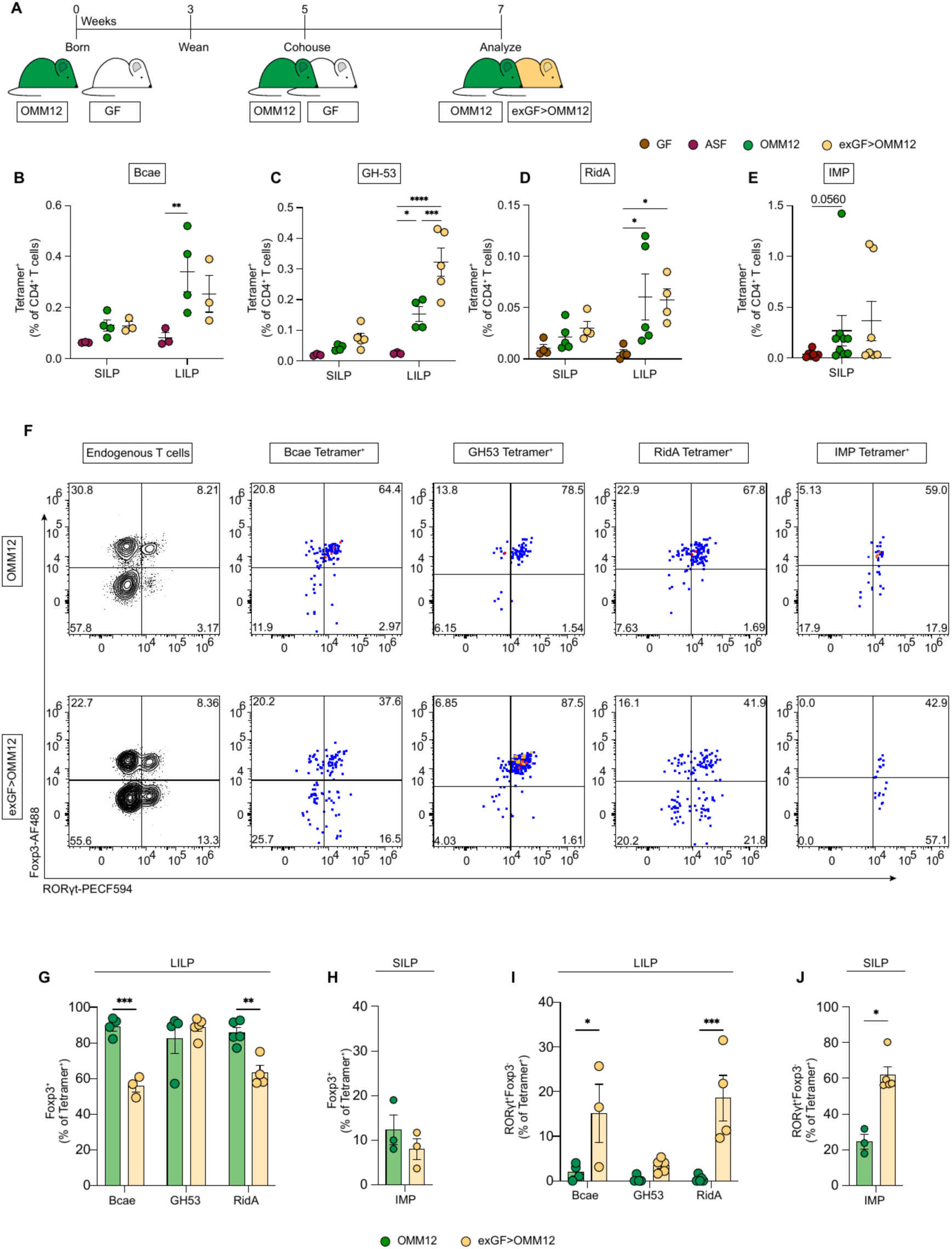
Horizontal colonization of germ-free mice with OMM12 partially uncouples microbiota-specific antigen recognition from stable Treg commitment. (A) Schematic of the experimental timeline. Briefly, OMM12-colonized breeder mice were mated to generate pups born into OMM12 (vertical colonization). At 5-7 weeks of age, germ-free (GF) mice were cohoused with OMM12-colonized mice for 2 weeks to permit horizontal colonization (exGF>OMM12), followed by analysis. (B-D) Summary plots of Bcae-, GH53-, and RidA-tetramer⁺ CD4⁺ T cells (% of CD4⁺ T cells) in the small intestinal (SILP) and large intestinal (LILP) lamina propria of GF, ASF, OMM12, or exGF>OMM12 mice. (E) Summary plot of IMP-tetramer⁺ CD4⁺ T cells (% of CD4⁺ T cells) in the small intestinal (SILP). (F) Representative flow cytometry plots of Foxp3 and RORγt expression among endogenous (tetramer-negative) CD4+ T cells and Bcae-, GH53-, RidA-, and IMP-tetramer+ CD4+ T cells (left to right) from the LILP of vertically colonized OMM12 (top) or horizontally colonized exGF>OMM12 (bottom) mice. Quadrant values indicate the percentage of cells within each respective parent gate (endogenous or tetramer+ CD4+ T cells). (G) Summary of Foxp3⁺ frequency (% of tetramer⁺ cells) among Bcae-, GH53-, and RidA-specific CD4⁺ T cells in the LILP (H) Frequency of of Foxp3⁺ frequency (% of tetramer⁺ cells) among IMP-specific CD4⁺ T cells in the SILP. (I) Summary of RORγt⁺Foxp3⁻ (Th17) frequency (% of tetramer⁺ cells) among Bcae-, GH53-, and RidA-specific CD4⁺ T cells in the LILP (J) Summary of RORγt⁺Foxp3⁻ (Th17) frequency (% of tetramer⁺ cells) among IMP-specific CD4⁺ T cells in the SILP. For (G) through (J), only mice with tetramer⁺ frequency exceeding the mean + 1 SD of the ASF control frequency were included (see Methods). Each dot represents an individual mouse; bars indicate mean ± SEM. For (B)-(D), (G) and (I) statistical significance was determined by two-way ANOVA with Sidak’s multiple comparisons test; *p < 0.05, **p < 0.01, ***p < 0.001. For (E), Statistical differences between groups were assessed using the Kruskal-Wallis test with Dunn’s multiple comparisons test. For (H) and (J), statistical differences were assessed using the Mann-Whitney test; *p < 0.05.

## DISCUSSION

By combining single-cell transcriptomic profiling with a scalable, single-cell-derived TCR reconstitution pipeline, we have characterized the endogenous CD4+ T cell response to an entire, defined intestinal microbiota with antigen-specific resolution during homeostasis and two distinct challenges to tolerance. Three important findings emerged. First, recognition of the microbiota at homeostasis was broad and antigen-diverse, spanning both private and shared epitopes across the entire community. Second, each species imprinted a distinct, largely reproducible T helper phenotype on its cognate T cells, yet the response was overwhelmingly Treg-biased when considered across the whole community. Third, when we used the resulting antigen-specific tetramers to track these same populations through two different perturbations of homeostasis, we found that the antigen-specific phenotype did not always follow the bulk compartment. Some antigens were remarkably stable while others were not, and this heterogeneity itself argues that the rules governing tolerance to the microbiota operate at the level of the individual antigen rather than uniformly across the population.

At homeostasis, recognition of the microbiota was essentially complete. Every member of the OMM12 consortium elicited at least one reactive TCR in our vertically colonized mice, arguing that the adaptive immune system samples the microbiota broadly rather than mounting responses to only a few immunodominant organisms, as has been described previously^4,8,9,62^. A system in which every antigen is culturable and therefore able to be screened likely explains why we observe broader recognition than has previously been appreciated. At the antigen level, this recognition was overwhelmingly private. Most reactive TCRs and the epitopes we identified for them were specific to a single species rather than shared across the community, in contrast to a recent report that found the opposite at the epitope level^13^. We did, however, identify a minority of cross-reactive epitopes, two of which (PGK and PfkB) map to highly conserved core metabolic enzymes, suggesting that cross-reactivity tracks how conserved a given peptide is across species rather than reflecting a promiscuous TCR recognition. Under this model, the shared antigen described elsewhere^13^ and our own PGK/PfkB/RidA epitopes likely reflect the same phenomenon of peptide conservation driving cross-reactivity.

Beyond this breadth, each species imprinted a stereotyped but distinct T helper phenotype on its cognate T cells, and not every response was a Treg. Amuc-reactive T cells were predominantly Tfh-skewed, agreeing with previous work from our group^13^, and Efae-reactive T cells were predominantly Th17-skewed, in line with evidence that Efae behaves as a pathobiont capable of driving colitis in IL-10-deficient mice^63^, and raising the possibility that Efae has features that bias T cells toward Th17 even without disease induction. Bani’s IMP epitope elicited a predominantly Th1 response, adding to a small list of bacterial species reported to elicit Th1 responses^18,64,65^. This diversity is notable given that non-Treg responses to the microbiota have typically been explained either as a property of a specific pathobiont or as a consequence of an already-inflamed environment, in which the antigen-specific repertoire constricts around a narrower, more pro-inflammatory set of specificities^22,23^. Even the recent report of a mixed Treg/Th17 response to individual gut bacteria^13^ was made in the context of horizontal colonization of adult mice. Here, Efae’s Th17 skew and Bani’s Th1 skew arise in vertically colonized mice under true homeostasis, indicating that this phenotypic diversity is not simply a byproduct of pathobiont status, active inflammation, or non-physiological colonization timing.

Despite this diversity, recognition at homeostasis was strikingly Treg-biased overall, with 11 of 12 OMM12 members eliciting at least one Treg-phenotype reactive TCR even when those same species also elicited other T helper subsets. Eclo, a Clostridia species, was a dominant inducer of Tregs, consistent with the well-established capacity of Clostridia to induce colonic Tregs^5,14^. Bcae, a Bacteroidetes species, elicited an almost purely Treg response and contributed the largest reactive population across both Bacteroidetes- and Firmicutes-reactive TCRs overall. This finding indicates that Treg-inducing capacity is not confined to Clostridia or Firmicutes, the phyla classically associated with Tregs^5,14,17,61,66^ but instead extends broadly across phyla. This aligns with prior work showing that Tregs make up the largest share of the CD4+ T cell response to the intestinal microbiota^66^, but extends this finding significantly. The homeostatic, vertically colonized response is genuinely diverse at the level of individual community members, yet overwhelmingly tolerogenic across the population.

Having developed the ability to track this diverse set of antigen-specific populations directly, we asked how they behaved under two very different perturbations of homeostasis: acute intestinal inflammation (piroxicam+αIL-10R) and colonization occurring outside the early-life window. Both perturbations produced a similar shift in the bulk CD4+ compartment, with a significant increase in Th17 cells in the intestinal LP, offering an opportunity to ask whether antigen-specific cells would behave the same way in similar environments. What we observed was surprising. The tetramer+ populations after piroxicam+αIL-10R challenge maintained the same phenotype as during homeostasis, but horizontal colonization resulted in altered phenotypes. Bcae and RidA, both predominantly inducers of Tregs at homeostasis, now induced Th17s upon horizontal colonization, and IMP-specific T cells, already predominantly Th1/Th17 at homeostasis, shifted still further toward Th17 induction. These findings may help explain why most prior work on intestinal Th17 responses to commensals, which involved new or altered colonization rather than mature tolerance^4,11,13,19,21^, reported outcomes different from what we observe in an already-established, vertically colonized response.

It is interesting to consider what distinguishes these two perturbations and causes such different changes in an antigen-specific manner, particularly because both perturbations caused similar expansion in the bulk Th17 populations in the intestines^67^. One possibility is that in piroxicam+αIL-10R-treated mice, inflammation is being induced on top of an already mature, homeostatic Treg population that may actively restrain a Th17 response to the same antigen, for instance by competing for access to cognate antigen-presenting cells (APCs). In horizontal colonization, cells are instead being primed for the first time, and the absence of a pre-existing cognate Treg response may allow for Th17 cells to be generated instead. This is consistent with a growing body of evidence that once a Treg population is stably established, it resists reprogramming even in an inflammatory environment that would divert a naive T cell encountering the same antigen for the first time^68^. This suggests that the population-level Th17 shift we observe with piroxicam+αIL-10R reflects recruitment of previously minor or newly activated specificities against additional, unmapped epitopes, rather than conversion of the Treg clones we characterized directly. A second possibility is that the antigen-presenting cell context matters. Building on foundational work showing that RORγt+ Treg populations can be induced by single species^61,69^, a recently described RORγt+ APC specializing in microbiota-reactive pTreg differentiation^70–74^ may only be available to, or only capture, certain antigens, making Treg commitment for those antigens more or less durable depending on how and where they are first presented. A third possibility is that Treg commitment for some antigens specifically depends on an early-life developmental window, while others, like GH53, are intrinsically tolerogenic regardless of the age or immunological context in which they are first encountered^75^. This could be because of the APC subset that predominantly captures and presents them^76^, their abundance, or their route of delivery^76,77^.

Critically, the three possibilities laid out above are not mutually exclusive. Not every antigen shifted under horizontal colonization, suggesting that the outcome of microbiota-specific T cells cannot be explained by the surrounding environment alone. If local cytokine and antigen-presenting cell context at activation were the dominant driver of CD4+ T cell fate, largely independent of antigen identity^74,78–80^, then antigens delivered within the same anatomical niche and time window should have converged on similar outcomes, yet GH53-specific cells remained stably Treg while Bcae-, RidA-, and IMP-specific cells did not. Our data instead argue for a model in which T cell identity is shaped, at least in part, by the identity of the antigen itself and its associated route of presentation, not solely by stochastic exposure to cytokines and other environmental cues encountered after activation.

Because our study for the first time comprehensively profiles the CD4+T cells response to the microbiota under homeostatic conditions, it may help reconcile many of the previous, apparently disparate findings related to recognition of the gut microbiota. Ultimately, this study illuminates that the rules governing microbiota-mediated CD4+ T cell responses are incredibly complex and far more diverse than a single organizing principle can capture.

## Limitations of the study

Our TCR screening was not exhaustive for both datasets and thus we may have missed some minor microbe-reactive clonotypes. The narrow filtering to focus on pTregs and Th17s in screening the inflammation dataset may have missed additional microbiota-reactive TCRs that were present in other clusters that were shown to have microbiota-reactive TCRs (e.g., Tfh). Despite validation of tetramer staining of hybridomas expressing cognate peptide TCRs, some tetramers did not stain meaningful populations of endogenous T cells.

## ACKNOWLEDGEMENTS

We thank Dr. S. Carroll for helpful discussions and comments on the manuscript; Dr. G. Victora for OMM12 ceca; Dr. C. Hsieh for the 58*ɑ*-β-hybridoma cell line; A. Liu, K. Siu, N. Butler, D. Levy, and H. Ahn for help with mice; L. Slayden for help with experiments; the NIH Tetramer Core at Emory University for production of MHC Ab tetramers; the UC Berkeley Cancer Research Lab Flow Cytometry Facility for assistance with flow cytometry; and QB3 Core Facilities for 10X library preparation and sequencing.

## AUTHOR CONTRIBUTIONS

Conceptualization, L.V., J.N.W., and G.M.B.; investigation, L.V., J.N.W., S.H.N, R.C.B, A.L.; formal analysis, L.V., J.N.W.; software development, J.N.W., A.T., A.W.; bioinformatic analysis, J.N.W.; writing – original draft, L.V.; writing – review and editing, L.V., J.N.W., R.C.B., G.M.B.; supervision, G.M.B.; funding acquisition, G.M.B.

## FUNDING

This work was supported by NIH grant AI200393 (to G.M.B.) and an Innovator Award from the Kenneth Rainin Foundation (to G.M.B.). L.V and J.N.W. were supported by an NIAID T32 Training Grant (T32AI100829).

G.M.B. is an Investigator of the Howard Hughes Medical Institute.

## CONFLICTS OF INTEREST

The authors declare no competing interests.

## DATA AND CODE AVAILABILITY

All data needed to evaluate the conclusions in the paper are present in the paper or supplemental information. Single cell data, including Seurat objects and all metadata, are deposited at GEO Accession GSE346897.

The primer design tools have been deposited at https://github.com/jwitch/SB-TCR-primer-design.

## MATERIALS AND METHODS

### Key Resources Table

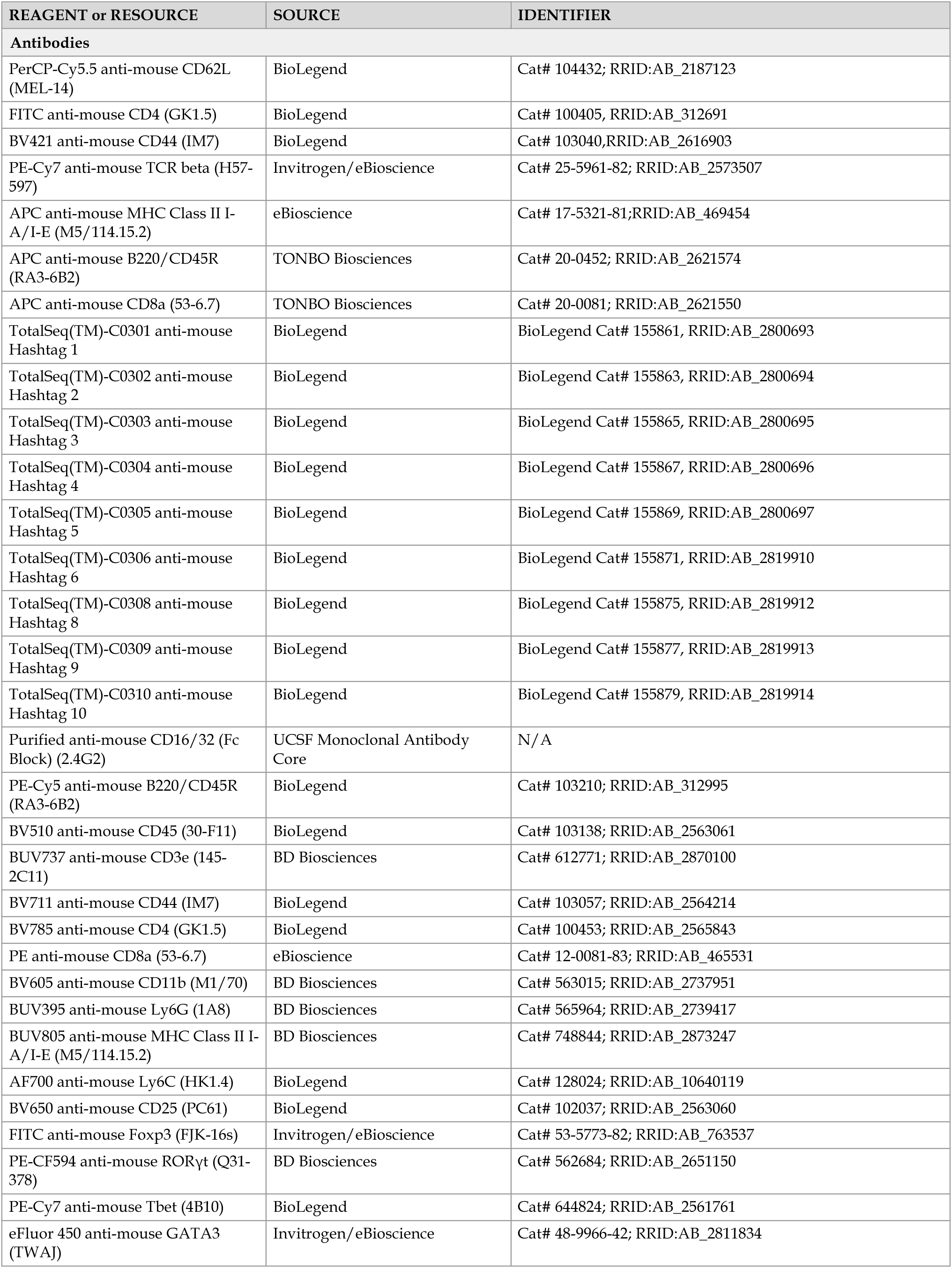
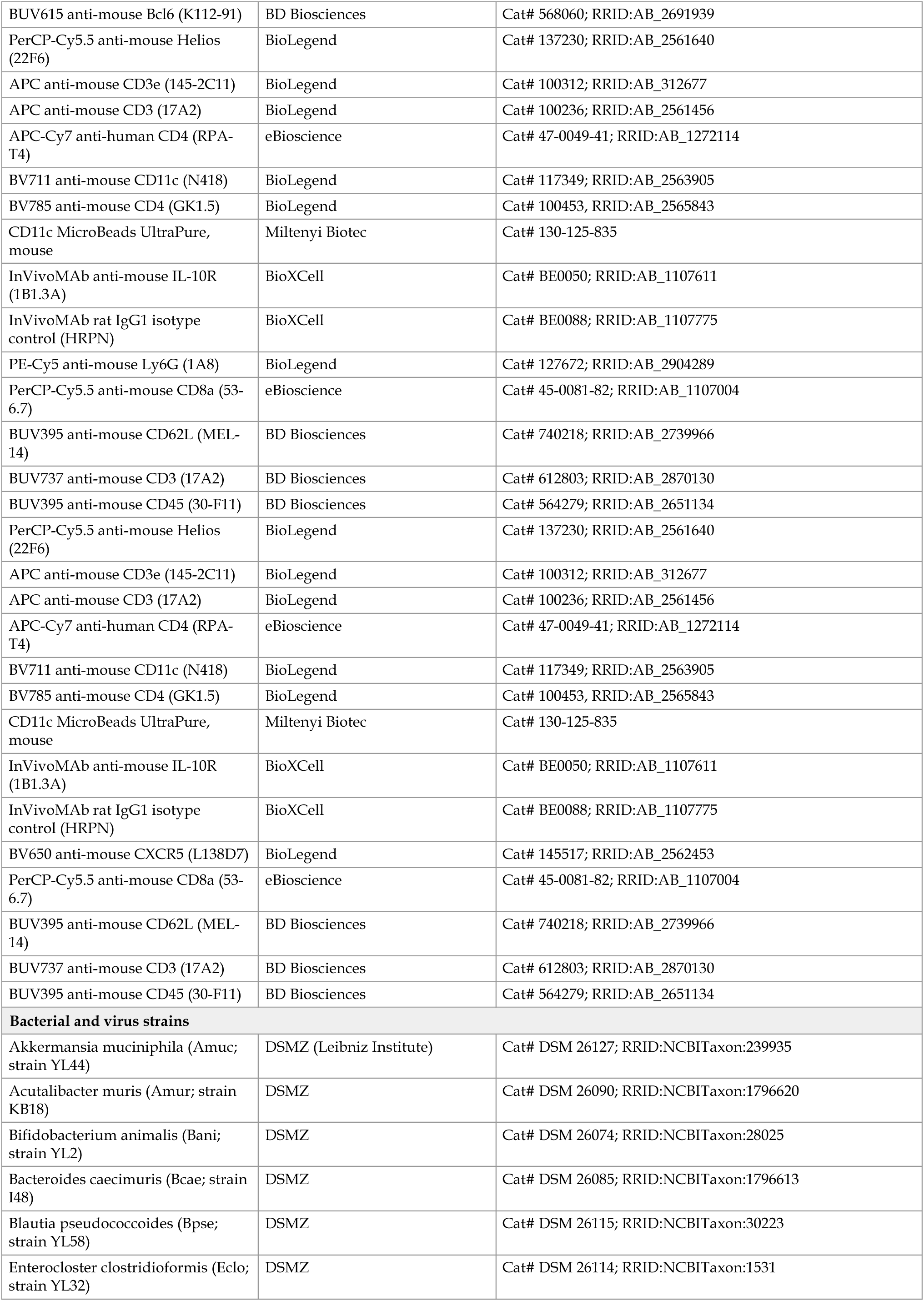
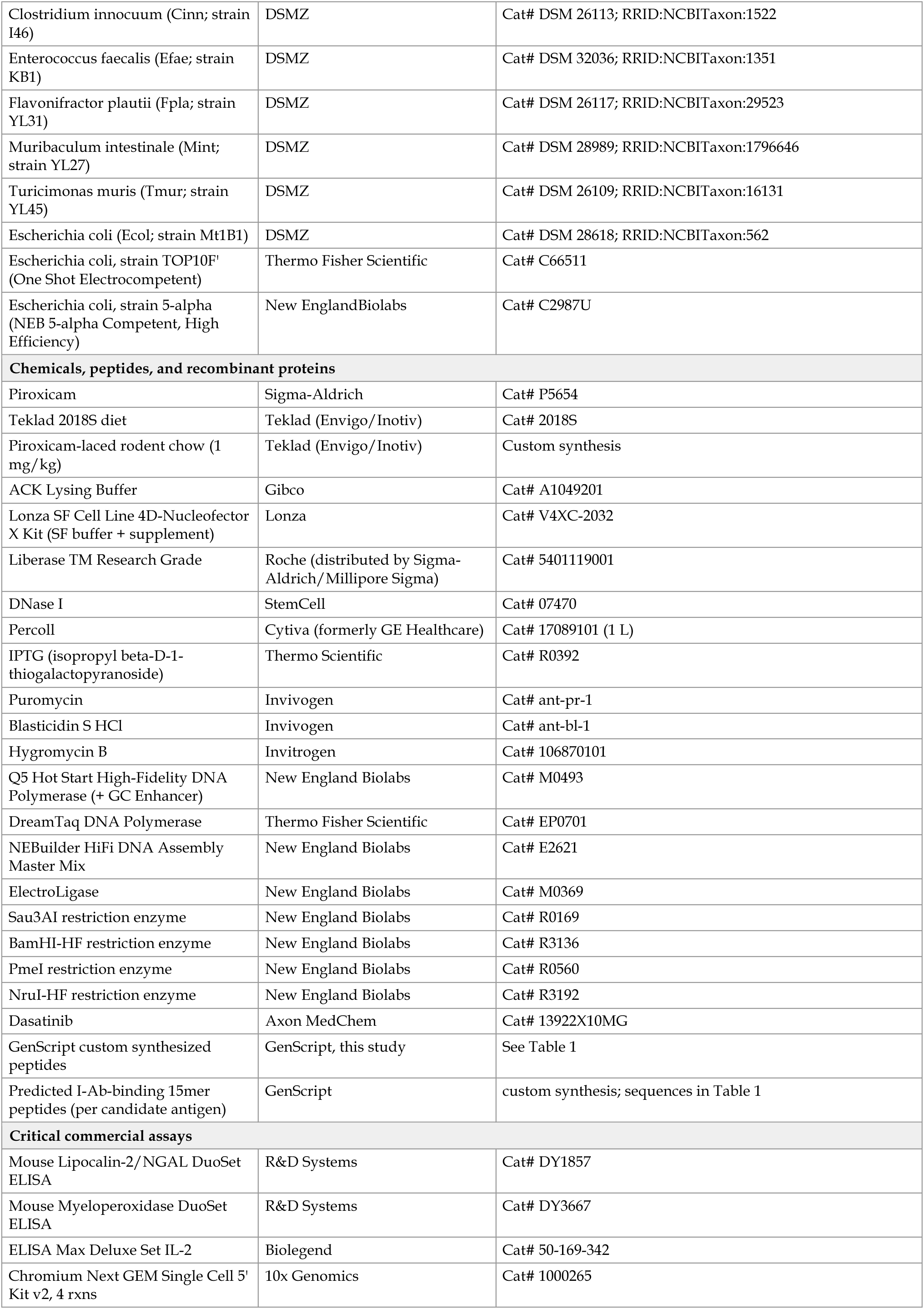
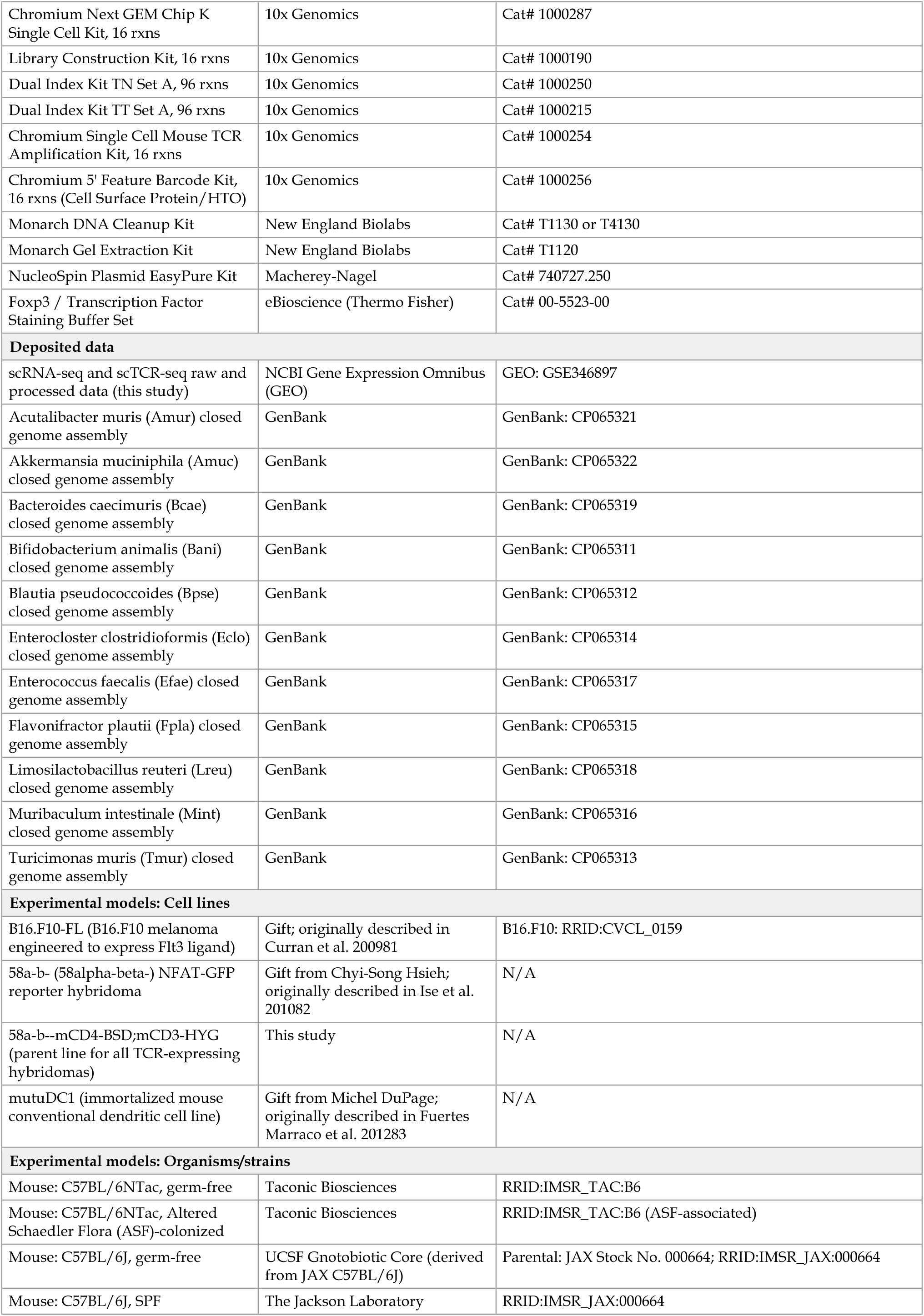
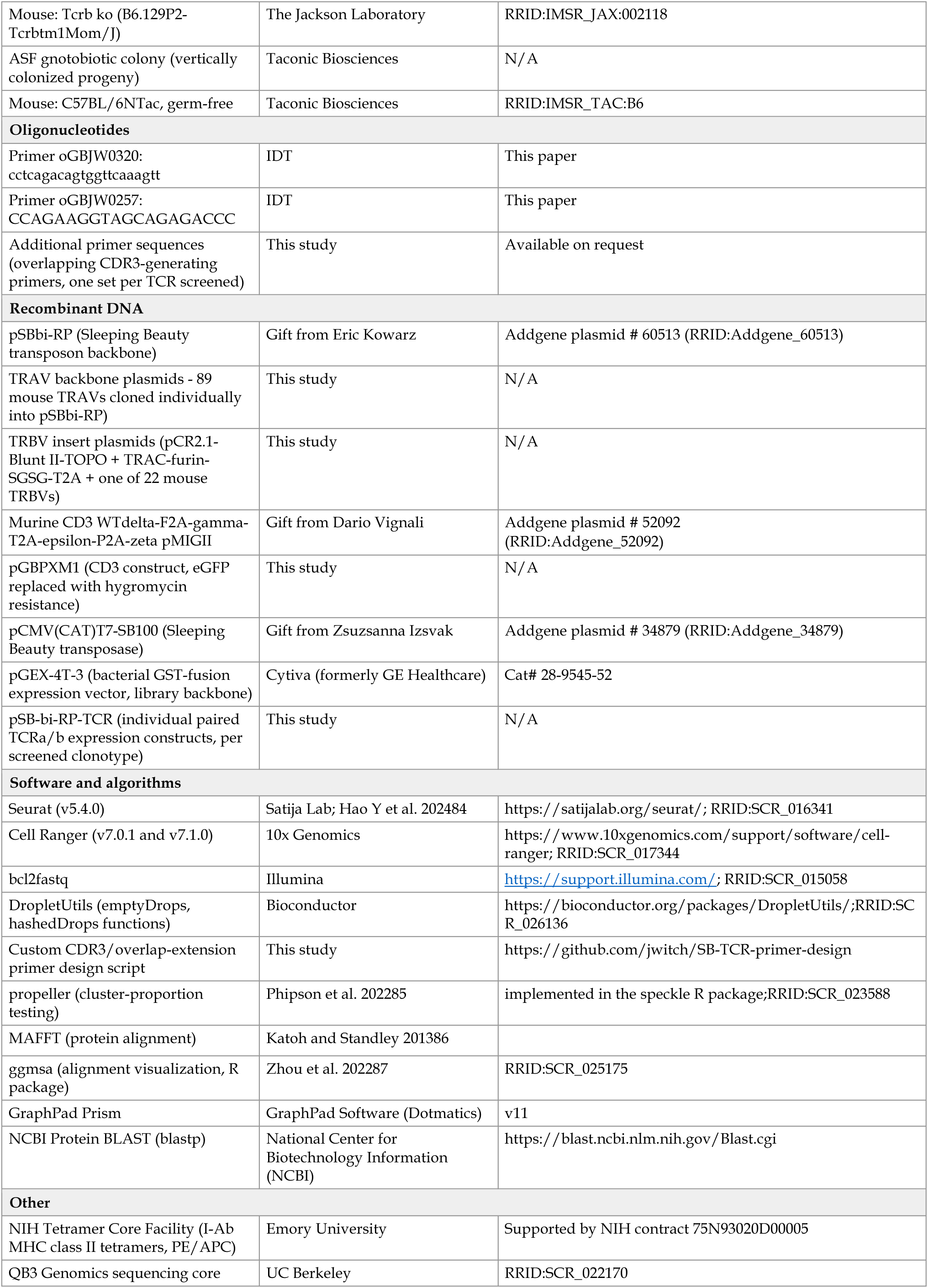
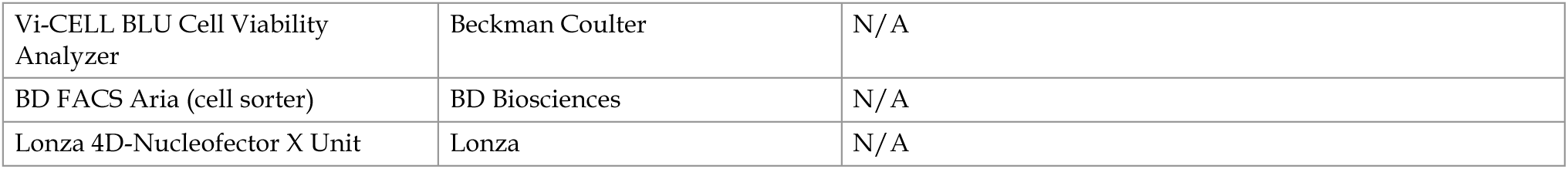

### Resource availability

**Lead contact** Requests for further information and resources should be directed to and will be fulfilled by lead contact, Gregory M. Barton.

**Materials Availability** Plasmids to generate TCRs, TCR expression plasmids or cell lines will be made available upon request via a materials transfer agreement with University of California Berkeley.

### Experimental model and study participant details

#### Mice

Mice were housed at the University of California, Berkeley under gnotobiotic or specific pathogen-free (SPF) conditions. All experiments were performed in accordance with an approved University of California Berkeley Animal Care and Use Committee protocol. Germ-free (GF) and Altered Schaedler Flora (ASF)-colonized C57Bl/6NTac (RRID:IMSR_TAC:B6) mice were obtained from Taconic Biosciences. GF C57Bl/6J (RRID:IMSR_JAX:000664) mice were obtained from the UCSF Gnotobiotic Core. All gnotobiotic mice were bred in flexible film isolators and used immediately upon removal or transferred to a Tecniplast Bioexclusion cage for experiments. C57Bl/6J and Tcrb-deficient (cat. no. 002118; RRID:IMSR_JAX:002118) mice were obtained from the Jackson Laboratory and used directly for experiments or bred in house under SPF conditions.

To generate an Oligo mouse microbiota (OMM)12 gnotobiotic mouse colony, male and female GF C5^7^Bl/6NTac mice housed in a flexible film isolator were gavaged with a slurry of luminal contents suspended in PBS from the cecum of an OMM12-colonized mouse that was a gift from Gabriel Victora at the Rockefeller University. Colonization of all 12 bacteria was confirmed by PCR and contamination ruled out by 16S sequencing. Harem mating cages were established from these mice and all subsequent experiments involving OMM12 mice were performed on vertically-colonized mice (progeny or their descendants) unless otherwise specified.

#### Acute inflammation model

6-8 week old male OMM12 colonized C57Bl/6NTac mice were moved from a flexible film isolator to an Tecniplast Bioexclusion cage. Mice were given 1 mg of αIL-10R (clone 1B1.3A, BioXCell cat. no. BE0050; RRID:AB_2894794) or isotype control (clone HRPN, Rat IgG1, κ, BioXCell cat. no. BE00^88^; RRID:AB_3667405) by intraperitoneal injection three days before starting mice on piroxicam-supplemented pellet mouse chow (px, Sigma-Aldrich cat no. P5654, 1 mg/kg, Teklad 2018S). A second αIL-10R injection was performed 7 days after the first injection (day 4 post-px). Mice were allowed to consume the chow ad libitum for 10 days. To check for intestinal distress, feces were collected and Lipocalin-2 (Lcn-2) was measured by ELISA (R&D DY1857). Males only were used in these experiments because the homeostasis 10x dataset was generated with males and females did not exhibit phenotypes as strong as the males.

#### Horizontal colonization model

GF mice, 5-7 weeks of age, were colonized with the OMM12 microbiota via bedding transfer from age-matched, OMM12-colonized donor mice. OMM12 donor bedding was refreshed every five days to establish horizontal transfer of the consortium, and intestinal T cell responses were subsequently analyzed as described above two weeks after initial bedding transfer (exGF>OMM12).

#### Bacterial strains

Pure, freeze-dried cultures were obtained from the Leibniz Institute German Collection of Microorganisms and Cell Cultures GmbH (DSMZ) for the following organisms: Akkermansia muciniphila (Amuc; YL44; DSM 26127; RRID:NCBITaxon:239935), Acutalibacter muris (Amur; KB18; DSM 26090; RRID:NCBITaxon:1796620), Bifidobacterium animalis (Bani; YL2; DSM 26074; RRID:NCBITaxon:28025), Bacteroides caecimuris (Bcae; I48; DSM 26085; RRID:NCBITaxon:1796613), Blautia pseudococcoides (Bpse; YL58; DSM 26115; RRID:NCBITaxon:30223), Enterocloster clostridioformis (Eclo; YL32; DSM 26114; RRID:NCBITaxon:1531), Clostridium innocuum (Cinn; I46; DSM 26113; RRID:NCBITaxon:1522), Enterococcus faecalis (Efae; KB1; DSM 32036; RRID:NCBITaxon:1351), Flavonifractor plautii (Fpla; YL31; DSM 26117; RRID:NCBITaxon:29523), Limosilactobacillus reuteri (Lreu; I49; DSM 32035; RRID:NCBITaxon:1597), Muribaculum intestinale (Mint; YL27; DSM 28989; RRID:NCBITaxon:1796646), Turicimonas muris (Tmur; YL45; DSM 26109; RRID:NCBITaxon:16131) and Escherichia coli (Ecol; Mt1B1, DSM 28618; RRID:NCBITaxon:562). For initial growth, strains were grown in media recommended by the DSMZ. Single use glycerol stocks for all strains were stored at −80°C for subsequent inoculations.

### Method details

#### Cell isolation from mouse tissues for T cell characterization

Mice were euthanized with CO2 inhalation followed by cervical dislocation. Spleens, mesenteric lymph nodes (MLN), small intestines (SI) and large intestines (LI, including cecum) were removed to ice cold media (RP-2 or RP-5; RPMI supplemented with 10 mM HEPES, 100 units/mL penicillin, 100 µg/mL streptomycin, 1x GlutaMAX, 2 or 5% fetal bovine serum and β-mercaptoethanol) for further processing as previously described7. Spleens and MLNs were mashed through a 70 µm strainer into a conical tube containing cold media. The cecal patch was removed and discarded from the cecum. The strainers were washed with media and cells were pelleted by centrifugation (500xg, 4°C, 5 min). MLN cell pellets were resuspended in 500 µl of media. To remove red blood cells, spleen cell pellets were resuspended in 1 mL ACK (Ammonium-Chloride-Potassium) lysis buffer (Gibco A1049201) and incubated for 3 min at room temperature (RT). Cells were washed with media, pelleted, and resuspended in a final volume of 5 mL of media. Peyer’s patches (PP) were clipped with curved scissors from the small intestine and mashed through a 100 µm strainer. The strainer was washed with media and PP cells were pelleted and resuspended in 500 µl of media.

For single cell experiments, small and large intestines were flushed with PBS and cut open longitudinally. Tissues were washed 3-6 times in PBS until liquid was clear. To strip epithelial cells from the small intestines, tissue was cut into 1-2 cm pieces and put into 15 mL of DTT solution (HBSS-10: HBSS without calcium/magnesium plus 10% FBS, 1x GlutaMAX, 10 mM HEPES, 100 units/mL penicillin, 100 µg/mL streptomycin; 1 mM dithiothreitol) and placed on a stir plate for 25 min at 37°C. The tissue was subsequently washed with PBS and transferred to 15 mL of EDTA solution (HBSS-0: HBSS without calcium/magnesium plus 1x GlutaMAX, 10 mM HEPES, 100 units/mL penicillin, 100 µg/mL streptomycin; 1.3 mM EDTA). Small intestine tissue was incubated for 20 min at 37°C with magnetic stirring. Epithelial cell removal from large intestines was achieved in one step by incubating in DTT+EDTA solution (HBSS-0; 1.3 mM EDTA; 1 mM DTT) with magnetic stirring at 37°C for 30 min. Intestinal tissues were washed with PBS over a 100 µm strainer and transferred to a 50 mL conical with PBS. Tissues were shaken in PBS about 20 times and poured over the 100 µM filter again. A final PBS wash was performed and tissue put into 15 mL of digestion buffer (RP-0; RPMI, 1x GlutaMAX, 10 mM HEPES, 100 units/mL penicillin, 100 µg/mL streptomycin, 5mg/mL collagenase VIII and 5 µg/mL DNase I) for 30 min to 1 hour depending on how long it took to digest the tissue to completion. Cells were filtered through a 100 µM strainer with any residual clumps being broken up with a syringe plunger and washed with 30 mL media. Immune cells were isolated at the interface of a 44/67% Percoll (Cytiva 17089101) gradient, washed with media and resuspended in 500 µL of media. For T cell phenotyping and tetramer experiments, small and large intestines were processed identically to remove epithelial cells in one step with DTT+EDTA solution for 35 min.

#### Sorting of antigen-experienced T cells from mice

For single cell sequencing experiments, isolated cells were subjected to a final 40 µm filtering step and stained with the following staining mix before sorting: LIVE/DEAD™ Fixable Near-IR (NIR; Invitrogen cat no. L10119, 1:1000), PerCP-Cy5.5 anti-mouse CD62L (clone MEL-14, Biolegend cat no. 104432; RRID:AB_2187123; 1:200), FITC anti-mouse CD4 (clone GK1.5, Biolegend cat. no. 100406, 1:400), BV421 anti-mouse CD44 (clone IM7, Biolegend cat. no. 103040, 1:400), PE-Cy7 anti-mouse TCRb (clone H57-597, Invitrogen 25-5961-82, 1:200), APC anti-mouse MHCII (clone M5/114.15.2, eBiosciences cat. no. 17-5321-81, 1:400), APC anti-mouse B220 (clone RA3-6B2, TONBO cat. no. 20-0452-U100, 1:400), APC anti-mouse CD8α (clone 53-6.7, TONBO cat. no. 20-0081-U100, 1:400). To distinguish cell origin during single cell analysis, a different Biolegend TotalSeqC hashtagging antibody (65.5 ng per million cells) was included in the surface staining for each sample. The entirety of each sample was run on a BD FACS Aria and cells sorted as CD44+CD62L-CD4+TCRb+MHCII-B220-CD8a-into 50% FBS in PBS. Cells were pelleted and resuspended in 100 µL of PBS. After counting on a hemacytomer, equivalent numbers of cells from each sample (MLN, PP, SILP and LILP from 2 mice) were mixed and pelleted. Cells were resuspended in the appropriate volume of PBS for subsequent single cell library synthesis.

#### Cellular phenotyping during inflammation

For immune cell profiling of mice treated with αIL10R and px, samples were first stained with NIR, and purified anti-mouse CD16/32 Fc Block (clone 2.4G2, UCSF core, 1:200). Then incubated with surface stain containing PE-Cy5 anti-mouse B220 (clone RA3-6B2, BioLegend cat no. 103210, 1:400), BV510 anti-mouse CD45 (clone 30-F11, BioLegend cat no. 103138, 1:200), BUV737 anti-mouse CD3ε (clone 145-2C11, BD cat no. 612771, 1:200), BV711 anti-mouse CD44 (clone IM7, BioLegend cat no. 103057, 1:200), BV785 anti-mouse CD4 (clone GK1.5, BioLegend cat no. 100453, 1:400), PE anti-mouse CD8α (clone 53-6.7, eBioscience cat no. 12-0081-83, 1:400), BV605 anti-mouse CD11b (clone M1/70, BD cat no. 563015, 1:1000), BUV395 anti-mouse Ly6G (clone 1A8, BD cat no. 565964, 1:200), BUV805 anti-mouse MHCII (clone M5/114.15.2, BD cat no. 748^84^4, 1:200), AF700 anti-mouse Ly6C (clone HK1.4, BioLegend cat no. 128024, 1:200), PE anti-mouse CXCR5 (clone L138D7, BioLegend cat no. 145504, 1:50), BV650 anti-mouse CD25 (clone PC61, BioLegend cat no. 102037, 1:200). Cells were then fixed and permeabilized using the eBioscience Foxp3/Transcription Factor Staining Buffer Set (Thermo Fisher cat no. 00-5523-00) and stained intracellularly with FITC anti-mouse Foxp3 (clone FJK-16s, Invitrogen cat no. 53-5773-82, 1:250), PE-CF594 anti-mouse RORγt (clone Q31-378, BD cat no. 562684, 1:500), PE-Cy7 anti-mouse Tbet (clone 4B10, BioLegend cat no. 644824, 1:200), eFluor 450 anti-mouse GATA3 (clone TWAJ, Invitrogen cat no. 48-9966-42, 1:50), BUV615 anti-mouse Bcl6 (clone K112-91, BD cat no. 568060, 1:100), and PerCP-Cy5.5 anti-mouse Helios (clone 22F6, BioLegend cat no. 137230, 1:100).

#### scRNA-seq and scTCR-seq library generation and analysis

Once sorted, cells were loaded onto 2 lanes of a Chromium Next GEM chip to achieve an intended recovery of 20,000 cells. Single cell cDNA was generated with the Chromium Next GEM Single Cell 5’ Kit v2 and Library Construction kit according to the manufacturer’s instructions (10x Genomics). For TCR sequencing, the Chromium Single Cell Mouse TCR Amplification Kit was used according to the manufacturer’s instructions. Hashtags in samples were detected using the 5’ Feature Barcode kit. Library intermediates and final sequencing libraries were analyzed using a Tapestation 4150.

Libraries were sequenced on a NovaSeq 6000 with Flow Cell S1 (Illumina) by QB3 Genomics (UC Berkeley, Berkeley, CA, RRID:SCR_022170) at a depth intended to capture ∼30,000 reads per cell for gene expression libraries and 5000 reads per cell for V(D)J and feature barcode libraries. Demultiplexing of libraries was achieved using bcl2fastq (Illumina). Fastq files were uploaded to the 10x Genomics Cloud and cellranger (version 7.0.1 or 7.1.0; RRID:SCR_017344)88 used to assign reads to transcripts (transcriptome mm10-2020-A), TCRs (vdj_GRCm38_alts_ensembl-7.0.0) or TotalSeqC barcodes.

For further analysis, a Seurat object84 (version 5.4.0, RRID:SCR_016341) was created for each of the homeostasis and inflammation datasets separately from the cellranger matrix files. Clonotype assignments were added as metadata. The TotalSeqC barcode library counts were added as a separate assay (“HTO”). The Seurat objects were then filtered to only include cells that had been assigned a TCR clonotype. The DropletUtils^89^ (RRID:SCR_026136) emptyDrops function was used to remove cells that had less than 500 feature library reads assigned to them. Then the hashedDrops function was used to assign cells to tissue samples. Any cells that were assigned as doublets by hashedDrops were removed. Additional filtering was done on cell quality (homeostasis: nFeature_RNA > 200 & nFeature_RNA < 6000 & nCount_RNA < 30000 & percent.mt < 5; inflammation: nFeature_RNA > 200 & nFeature_RNA < 7000 & nCount_RNA < 60000 & percent.mt < 5). For individual datasets, the seurat pipeline with the functions NormalizeData, FindVariableFeatures, ScaleData, and RunPCA was used for data processing. For clustering the homeostasis dataset, FindNeighbors was run with 20 dimensions and FindClusters with a resolution of 0.7. For the combined dataset analysis, the homeostasis and inflammation Seurat objects were merged after RunPCA,integrated using CCAIntegration, and layers rejoined on the RNA assay. FindNeighbors was run with 20 dimensions and FindClusters with a resolution of 0.6 were used for clustering. Marker genes used to identify each annotated cluster in the homeostasis-only dataset were as follows: GALT central memory thymic (t)Tregs - Foxp3+Ikzf2+Nrp1+Gpr83+Il2ra+Lrrc32+Itgae+, LP effector (e)Tregs - Foxp3+Rorc-Il1rl1(ST2)+Klrg1+Areg+Gata3+Rora+Ccr8+Ctla4+Icos+Lrrc32+Itgae+, LP peripheral (p)Tregs - Foxp3+Rorc+Il10+Ctla4+Havcr2+Maf+Lag3+ (and majority SILP and LILP barcodes), GALT pTregs - Foxp3+Lag3+Icos+Tigit+Maf+Ccr4+Tnfrsf4/18+ (and majority MLN and PP barcodes), circulating, recently- stimulated Th1 - Tbx21+Junb+Fos+Cxcr3+Gzma+Ifng+Ccr9loCd40lg+, gut-resident cytotoxic Th1 - Il12rb2hiCxcr6+Gzmk+Gzmb+Tbx21+Ifng+Ccr9hi, Th2 -Gata3+Il4+Il5+, Th17 - Rorc+Foxp3- Il17a+Il17f+Il23r+Ccr6+, pre-T follicular helper (fh) -Tcf7+Slamf6+Id3+Cd200+Lck+ (and PP tissue identity), Tfh - Bcl6+Cxcr5+Pdcd1+Tox2+Il21+, naive (or T stem-like central memory, Tscm) - Ccr7+Sell+Cd44- Lef1+Satb1+S1pr1+, recently activated, uncommitted (Tact) - S1pr1+Klf2+Rasgrp2+Itga4+, blasting - Myc+Srm+Bcat1+nucleolar/ribosome-biogenesis genes, proliferating -Mki67+Top2a+Ccnb1/2+Cdk1+Birc5+, gamma delta (gd) T (1) - Tcrg-V4+Trgv2+Tcrg-C1/C2/C4+Zbtb16+Klrb1c+, gd T (2) - Tcrg-C4+Trgv2+Tcrg- C1/C2+Zbtb16+Klrb1c+, interferon-stimulated genes (ISG)-high - Isg15+Ifit1/3+Irf7+Stat1/2+Rsad2+. A dotplot of a reduced set of these genes that distinguish individual clusters is reported in Figure S1. gd T (1) and gd T (2) are likely contaminating populations of cells not removed during FACS but inclusion did not affect the interpretation of other Th phenotypes so further filtering of the data was not performed.

Marker genes used to identify each annotated cluster in the combined dataset were as follows: GALT central memory thymic tTregs - Foxp3+Ikzf2+Nrp1+Gpr83+Cd27+, LP eTregs -Foxp3+Klrg1+Areg+Ccr8+Tnfrsf9+Tnfrsf4+Tigit+Gata3+, LP pTregs - Foxp3+Rorc+Il10+Ctla4+Havcr2+Maf+Lag3+, GALT pTregs -Foxp3+Lag3+Icos+Maf+Il2ra+Rorc(moderate), circulating, recently-stimulated Th1 -Tbx21+Junb+Fos+Fosb+Jun+Nr4a1/2+Cxcr3+Ifng+Il12rb2lo, gut-resident cytotoxic Th1 -Il12rb2hi+Ccr9hi+Gzma+Ifng+Tbx21+Nkg7+Xcl1+, Th2 - Il4+Gata3+Il17rb+Il2+, Th17 -Il17a+Il17f+Il22+Il23r+Rorc+Cxcr6+, pre-Tfh - Tcf7+Slamf6+Id3+Themis+, Tfh - Cxcr5+Pdcd1+Bcl6+Tox2+Il21+, naive (or Tscm) - Ccr7+Sell+Lef1+, GALT recently activated - S1pr1+Klf2+Rasgrp2+Itga4+Il7r+, LP recently activated -Rel+Nfkb1+Nfkbiz+Nr4a2/3+Gadd45b+, blasting - Srm+Nop16+Nhp2+Gar1+Fbl+ (nucleolar/ribosome-biogenesis genes), proliferating - Mki67+Top2a+Birc5+Mcm2-7+Rrm1/2+, Cytotoxic effector - Gzma+Gzmk+Nkg7+Ccl5+Ctsw+Cd160+, gd T (1) - Tcrg-V4+Trgv2+Tcrg-C1+Zbtb16+Klrb1c+, gd T (2) - Tcrg-C4+Trgv2+Tcrg-C1/C2+Zbtb16+Klrb1c+. A dotplot of a reduced set of these genes that distinguish individual clusters is reported in Figure S2.

For broader clustering of T cell phenotypes used in the main text, the Seurat clusters that went into each category are as follows: Tregs - GALT central memory tTregs, LP eTregs, LP pTregs, GALT pTreg; Th1 - circulating, recently-stimulated Th1, gut-resident cytotoxic Th1; pre-Tfh -pre-Tfh; Th2 - Th2; Tfh - Tfh; naive - naive (or Tscm); recently activated - recently activated, uncommitted, GALT recently activated, LP recently activated, blasting; other - proliferating, ISG-high, cytotoxic effector, gd T(1), gd T (2).

Expanded T cell clonotypes were defined as any clonotype that had at least 3 cells within a cluster. For homeostasis derived samples, clonotypes that were chosen to be screened included the top 50 most abundant clonotypes and any clonotype in clusters Th2, Th17, LP pTregs, LP pTregs, Tfh and pre-Tfh where the within cluster cell number was ≥ 3 cells. For inflammation datasets, expanded clonotypes were chosen to test for reactivity from the top 50 most abundant clonotypes and for clonotypes in the LP pTreg (the highest microbiota reactivity was found in this cluster during homeostasis) and Th17 (frequency of cells significantly expanded during inflammation) clusters defined from the combined data analysis for which there were ≥ 3 cells within the cluster.

To test for T cell cluster proportion shifts between homeostatic (homeo) and inflammatory (infl) conditions, we used propeller^85^, implemented in the speckle R package (RRID:SCR_023588), which tests for differences in cell-type proportions between conditions using biological replicates rather than treating individual cells as the unit of replication. Each condition (homeo, infl) comprised two mice, with tissue and animal identity assigned to each cell from its cell-hashing classification (Hashing_Classification), cross-checked against the hashtag oligo identity (HTO_maxID) to remove rare mis-hashed cells; only cells classified as doublets were excluded. Because each mouse contributed one sample per gut tissue (MLN, PP, SILP, LILP), propeller was run separately within each tissue (n = 2 vs. n = 2 biological replicates per test) rather than pooling across tissues, at two levels of cluster resolution: a fine resolution corresponding to the full set of seurat clusters, and a broad resolution corresponding to nine collapsed phenotypic categories (naive, Treg, Th1, Th2, Th1^7^, pre-Tfh, Tfh, recently activated, and other). Cell-type proportions were arcsine–square-root transformed and compared using propeller’s empirical Bayes moderated t-test, with clusters representing fewer than 10 cells in any individual sample excluded from a given tissue-by-resolution comparison prior to variance estimation (independent filtering), to prevent proportion estimates derived from very small cell numbers from contributing noise to the moderated variance shared across better-powered clusters. P-values were corrected for multiple testing within each tissue-by-resolution comparison using the Benjamini-Hochberg procedure, and clusters were considered significantly expanded or depleted during inflammation at FDR < 0.05.

Graphics showing single cell T cell phenotype quantifications were generated by adding reactivity metadata (Table S2) for all TCRs tested to the Seurat object metadata and then using a custom script to pull and quantify the number of cells and TCRs for a specific reactivity.

#### Antigen presenting cell isolation from mice for TCR hybridoma coculture assays

Tcrb-deficient mice (6-16 weeks old) were injected subcutaneously into the rear flank with 1x107 B16.F10 cancer cells expressing Flt3 ligand (B16.F10-FL)^81^. 7-14 days after injection, tumor-bearing mice were euthanized and spleens harvested. 2- 3 spleens were placed in 10 mL prewarmed digestion solution (HBSS containing magnesium and calcium (HBSS+) plus liberase TM and DNase I). An insulin syringe was used to inject the spleens with 100-200 µL of digestion solution. Spleens were incubated for 20 min at 37°C and then minced. Minced spleens were incubated for a further 10 min at 37°C then mashed through a 70 µm strainer. The strainer was washed with 20 mL RP-5 to quench the reaction and cells pelleted. Media was removed and the pellet was resuspended in 3 mL ACK for 5 min. 20 mL of media was added and cells pelleted. Cells were resuspended in RP-10+ and counted on a Vi-CELL BLU instrument (Beckman Coulter). CD11c+ antigen-presenting cells were enriched using CD11c MicroBeads UltraPure (mouse) (Miltenyi Biotec, no. 130–125-835) according to manufacturer’s instructions.

Purified APCs were counted and resuspended to 1.5x10^6^ cells/mL in RP-10+. 100 µL was transferred to each well of a 96 well round bottom TC-treated plate to be used the same day in the TCR stimulation assay.

#### TCR cloning strategy

The mouse TCR cloning strategy was inspired by the human TCR cloning system developed previously^43^. Briefly, a Sleeping Beauty transposon is constructed that contains a paired TCRα and TCRβ chain derived from single cell sequencing. The Sleeping Beauty transposon plasmid, pSBbi-RP, was a gift from Eric Kowarz (Addgene plasmid #60513; http://n2t.net/addgene:60513; RRID:Addgene 60513). One of 89 mouse TRAVs, a PmeI cut site, a NruI cut site, and TRBC1 were cloned into pSBbi-RP to create plasmids that serve as the backbone for TCR cloning (TRAV plasmid; sequences can be found in Table S1). TRAV plasmids were prepared for cloning by restriction digest with PmeI and NruI-HF and DNA cleanup with a Monarch DNA cleanup kit (NEB T1130 or T4130). To create plasmids to serve as TRBV inserts, a PmeI cut site, the TRAC sequence, a furin-SGSG-T2A cleavage sequence, one of 22 mouse TRBVs, and a second PmeI cut site were cloned into the pCR2-Blunt II-TOPO vector (TRBV plasmid; sequences can be found in Table S1). Prior to cloning, the TRBV plasmids were cut with PmeI and the 0.8 kb insert was gel extracted with a Monarch Gel Extraction kit (NEB T1120). Further purification before Gibson assembly was performed using a Monarch DNA cleanup kit.

Overlapping primers were designed (by hand or using a custom script deposited on Github) to generate the missing CDR3 sequences with homology to the variable and constant regions of their respective chain by PCR. Q5 Hotstart polymerase with GC enhancer was used to generate the PCR products with the following cycling conditions: 98°C 30 s, 5 cycles of 98°C 5 s, 55°C 10 s, 72°C 5 s, 25 cycles of 98°C 10 s, 72°C 5 s, and a final extension of 30 s at 72°C. CDR3 PCR products were purified using a Monarch DNA kit and resuspended to 10 ng/µL for downstream use. To create the insert to the TRAV plasmid, CDR3α PCR product, CDR3β PCR product and TRAC-TRBV fragment were used in an overlap extension PCR reaction as previously described.^90^ Following amplification, PCR products were run on a gel to check for multiple bands and purified using a Monarch DNA cleanup kit (single band) or, if multiple bands were present, the correct size band was extracted. For assembly, a 10 µL DNA solution was made containing 200 ng of TRAV plasmid and 100 ng of CDR-TRBV insert in water. DNA solution was mixed 1:1 with NEBuilder HiFi DNA Assembly Master Mix and incubated at 50°C for 15 min.

Assembly reaction (0.5 µL) was transformed into 5-alpha competent cells (NEB C2987U, 25 µL) following manufacturer’s instructions except for recovery in 150 µL of Super Optimal Catabolite (SOC) medium. 5 µL of transformed cells was plated to LB+carbenicillin (carb) and incubated overnight at 37°C. To check for insertion, the next day, 3-8 transformants per transformation were individually resuspended in 12 µL of 2x Yeast Extract Tryptone (2xYT)+carb in a PCR tube strip or plate. Colony PCR was performed with DreamTaq polymerase, 1 µL of culture, and primers oGBJW0320 and oGBJW0257 using the following cycling conditions: 95°C 10 min, 35 cycles of 95°C 30 s, 53°C 30 s, 72°C 1.5 min, and a final extension of 10 min at 72°C. For transformants with 1.5 kb inserts, 5 mL overnight cultures (2xYT+carb) were inoculated with 5 µL of resuspended transformants. Plasmids were purified using the Nucleospin Plasmid EasyPure kit (Macherey-Nagel cat no. 740727.250) using twice the standard volumes of A1, A2, and A3 and resuspending in 30 µL ultrapure water. Purified plasmids were sent for sequencing with oGBJW0320 and oGBJW0257. If the miniprep concentration was greater than 500 ng/µL, the plasmid was used directly in cell line generation. If not, the miniprep was concentrated by precipitation to reduce the volume required during nucleofection. A complete list of all TCR sequences cloned can be found in Table S2.

#### TCR-expressing cell line generation

The NFAT-GFP reporter mouse 58*ɑ*-β- hybridoma cell line^82^ was a gift from Chyi-Song Hsieh. Human CD4 was originally used to select for reporter integration. Stable expression of mouse CD4 in this cell line was established via transduction of a plasmid (pGBPXM2-1b) containing mouse CD4 and blasticidin resistance (58*ɑ*-β-mCD4-BSD). After selection with blasticidin (25 µg/mL), CD4 expression was confirmed by flow cytometry. Murine CD3 WTdelta-F2A-gamma-T2A-epsilon-P2A-zeta pMIG II was a gift from Dario Vignali (Addgene plasmid # 52092; http://n2t.net/addgene:52092; RRID: Addgene_52092). This plasmid was modified to replace eGFP with hygromycin resistance (pGBPXM1). After transduction into 58*ɑ*-β-mCD4-BSD, cells were selected in 50 µg/mL hygromycin (58*ɑ*-β-mCD4-BSD;mCD3-HYG). To confirm expression of CD3 by flow required introducing a TCR to the cell line, which was achieved using the Am124 TCR. 58*ɑ*-β-mCD4-BSD;mCD3-HYG served as the parent cell line for all TCR-expressing hybridomas that were used in analysis of reactivity.

To introduce TCRs into the 58*ɑ*-β-mCD4-BSD;mCD3-HYG cell line, 1x10^6^ cells were resuspended in 20 µL SF buffer with supplement (Lonza V4XC-2032) plus 0.75 µg pSB-bi-RP-TCR plasmid and 0.25 µg pCMV(CAT)T7-SB100 (a gift from Zsuzsanna Izsvak; Addgene plasmid # 34879; http://n2t.net/addgene:34879; RRID:Addgene_34879) in a 16 well cassette. Program CM-150 on a 4D Nucleofector X Unit (Lonza) was used for transfection. 80 µL of RP-10 was added to the well and incubated for 5 min at RT. The total volume in the well was transferred to a 96 well non-treated culture plate with 150 µL of prewarmed RP-10 and incubated overnight at 37°C with 5% CO2. The following day, the whole volume from the 96 well plate was transferred to 1 mL of media in a 24 well plate and incubated overnight. To check transfection efficiency, the leftover cells were stained with stimulation stain mix: LIVE/DEAD™ Aqua (Aqua, Invitrogen cat. no. L34957; 1:1000), APC anti-mouse CD3ε (clone 145-2C11, Biolegend cat. no. 100312, 1:200) or APC anti-mouse CD3 (clone 17A2, Biolegend cat. no. 100236, 1:200), APC-Cy7 anti-human CD4 (clone RPA-T4, eBioscience cat. no. 47-0049-41, 1:200), PE-Cy7 anti-mouse TCRb (clone H57-597, Invitrogen 25-5961-82, 1:200), BV711 anti-mouse CD11c (clone N418, Biolegend 117349, 1:500), and BV785 anti-mouse CD4 (clone GK1.5, TONBO cat. no. 20-0041-U100, 1:400). The next day, 1 mL of culture was spun down and resuspended in 4 mL RP-10+ with 2 µg/mL puromycin in a 6 well culture dish. After 2 days, media was replaced with fresh RP-10+ with puromycin. After 2 more days, 200 µL of selected cells was stained with stimulation staining mix to check selection efficiency and TCRb expression. If TCRb+, each cell line was either used immediately for stimulation assay or cryopreserved. Occasionally, despite the correct TCR sequence from single cell data in the plasmid, there was no expression of TCRβ and those strains were not pursued further (classified as NE, not expressed in metadata).

#### Antigen preparation for TCR stimulation assays

Amur, Bani, Bpse, Eclo, Cinn, Efae, Fpla and Ecol were grown in Anaerobic Akkermansia Media (AAM: 18.5 g/L Brain Heart Infusion, 5 g/L yeast extract, 15 g/L trypticase soy broth, 2.5 g/L K2HPO4, 0.5 g/L glucose, 0.4 g/L Na2CO3, 0.5 g/L cysteine-HCl, 1 µg/mL resazurin, 5 mg/L menadione, 3% fetal bovine serum and 1 mg/L hemin) at 37°C in a flexible film anaerobic chamber (COY laboratories) until saturation. Amuc was grown in AAM supplemented with 2.5 g/L hog gastric mucin or Akk synthetic media (0.4 g/L KH2PO4, 0.53 g/L Na2HPO4, 0.3 g/L NH4Cl, 0.3 g/L NaCl, 0.1 g/L MgCl2·6H2O, 0.15 g/L CaCl2· 2H2O, 0.5 mg/L resazurin, 4 g/L NaHCO3, 0.45 g/L L-cysteine·HCl, 3 mL trace mineral solution (ATCC – TMS), 16g/L soy peptone, 4g/L L-threonine, 5.5g/L N-acetylglucosamine, 4.5 g/L glucose) under anaerobic conditions. Bcae was grown in AAM supplemented with fetal bovine serum up to 10% or Chopped Meat Carbohydrate Broth (BD 288130) under anaerobic conditions. Mint was grown in Chopped Meat Carbohydrate Broth under anaerobic conditions. L. reuteri was grown in de Man, Rogosa, and Sharpe (MRS) broth (BD 288210) under microaerophilic conditions. Tmur was cultured as a lawn on Colombia agar (Sigma-Aldrich 27688-100G) with 10% defibrinated sheep’s blood (Thomas Scientific C838N51) at 37°C in an anaerobic chamber for 2-3 days. For liquid cultures, at saturation, OD600 readings were taken and bacteria were pelleted for 30 min at 4000xg in a 4°C table top centrifuge. For Tmur plates, cells were scraped off into PBS and vortexed to create a single cell suspension from which the OD600 was taken then the suspension was pelleted. Cells were resuspended at an OD600 of 10 in sterile PBS and stored at −80°C in screw cap vials. To heat-kill the bacteria, a vial was thawed and 100 µL aliquots into PCR tubes. The tubes were heated to 95°C for 30 min in a thermocycler then placed back at −80°C until used in stimulation assays.

For in vivo derived antigens, contents from the ceca of GF or OMM12 colonized mice was resuspended to 1 g/mL in PBS and subjected to a low g (200xg) spin for 5 min. Supernatant was removed to a new tube and subjected to a high g (>10,000xg) spin for 5 min. The supernatant was removed, the volume of the supernatant was measured and the pellet resuspended in the same volume. The resuspended pellet was heat-killed as above.

#### TCR stimulation assay

For initial screening of reactivity, 2.5x10^4^ TCR-expressing cells were incubated with 7.5x10^4^ CD11c+ antigen-presenting cells, and 3.^7^5x107 heat-killed bacteria from monocultures (MOI=500) or heat-treated luminal contents from GF or OMM12 colonized mice in a total volume of 200 µL RP-10+. As a positive control for TCR stimulation, a final concentration of 1 µg/mL αCD3ε was added to cocultures. Hybridomas incubated with or without APCs only were used as negative controls. After an overnight incubation at 37°C with ^5^% CO2 (1^4^-16 h), plates were spun down and cells stained with stimulation stain mix. GFP-expression in huCD4+CD11c-cells was measured by flow cytometry. Heatmaps showing reactivity are reported as log2((%GFP upon stimulation-%GFP of hybridoma only)/(%GFP of αCD3ε control-%GFP of hybridoma only)). Positive reactivity was determined as >2. For weak hits, IL-2 was measured in a coculture assay with mutuDC1s (23162549; a gift from Michel DuPage) and heat killed bacteria. If the reactivity did not validate, the TCRs were excluded from additional analyses.

For antigen screening, 1x105 of mutuDC1 cells were preplated in 100 µL of mutuDC media (IMDM supplemented with 10% FBS, 1 mM HEPES, pen/strep, GlutaMAX and Bme) the day before stimulation. 4 to 5 TCR-expressing cell lines were mixed together in mutuDC media and added to the adhered mutuDC1s such that 1.25x104 of each TCR hybridoma were present per well. To each well, 2.5 µL of heat-killed expression library bacteria was added and plates incubated for 24 h at 37°C with 5% CO2. 150 µL of culture supernatant was transferred to a 384 deep well plate and stored at −80°C until thawed to measure IL-2. 30 µL was used to perform an IL-2 ELISA in 384 well format.

#### Bacterial expression library for antigen discovery

Bacterial protein expression libraries were synthesized as previously described with minor modifications^7,13^. Genomic DNA (gDNA) was isolated from monocultures of Amur, Bani, Bcae, Eclo, and Efae using the DNeasy PowerSoil Pro Kit (Qiagen). 40-100 µg of gDNA was subjected to partial digestion with Sau3AI. The volume of SauAI added to the first reaction was the amount required to fully digest the amount of gDNA present. SauAI was serially diluted 1:2 and the same amount of gDNA was added to each reaction. Reactions were allowed to proceed for 10 min at 37°C and then heat-inactivated. Digests were pooled and fragments from 0.5-5 kb were gel extracted. These fragments were ligated into BamHI-HF-digested pGEX-4T3 with ElectroLigase. Five µL of the ligation reaction was electroporated into 100 µL of TOP10F’ Electrocompetent cells (Thermo Fisher cat. no. C66511). After 1 hour of recovery in SOC, 10 µL of cells were plated to LB+carb and incubated overnight at 37°C while the rest were left in SOC at 4°C. Colony forming units were counted to estimate transformation efficiency. The rest of the recovered cells were resuspended to 30 colonies per 150 µL of 2xYT+carb in 384 deep well plates. Culture plates were shaken overnight at 37°C. 50 µL of culture was transferred to another 384 well plate containing 25 µL of 50% glycerol and stored at −80°C. 20 µL of culture was transferred to a third 384 well plate containing 180 µL 2xYT+carb and shaken for 2-3 h until OD600 ∼0.7-1. Protein expression was induced by adding 5 µL of 40 mM IPTG and incubating at RT overnight with shaking. Bacteria were pelleted by centrifuging the plates at maximum speed for 30 min at 4°C in a bucket centrifuge. Media was decanted and cell pellets were resuspended in 20 µL of PBS and transferred to a 384 well PCR plate. Bacteria were heat-killed at 95°C for 30 min in a thermocycler. Plates were stored at −80°C until used in stimulation assays.

Once stimulation assays were performed and supernatants assessed for IL-2 by ELISA, positive library wells were retested against individual TCRs from the mixed hybridoma pool to determine which TCRs reacted to the antigen pool. The glycerol stock corresponding to any well that was positive for IL-2 was streaked for singles and individual colonies prepared for stimulation assays as described above. Individual colonies were tested against reactive TCRs to narrow the reactivity to a single clone. Individual bacterial strains that induced IL-2 were grown up for plasmid isolation. Plasmids or bacteria clones were then sent for whole plasmid sequencing at Plasmidsaurus. The genomic DNA insert was evaluated for the presence of open reading frames. The amino acid sequence of any gene fragment found, regardless of whether it was in frame with glutathione S-transferase (GST) in the expression plasmid, was evaluated for epitopes that were predicted to bind the mouse class II I-Ab allele using NetMHCIIpan-4.3^91^ (RRID:SCR_025434). From the 9mer peptide cores that had a peptide that ranked as a strong binder (rank <1%), the 15mer with the lowest rank value encompassing that core was ordered for testing reactivity (GenScript, crude purity). If there were no strong binders, the predicted strongest binding 15mers were ordered that encompassed weak (rank <5%) binding cores. A complete list of identified peptides can be found in Table 1.

#### Peptide alignments

EcloPfkB and BaniPGK were used in Protein BLAST (https://blast.ncbi.nlm.nih.gov/Blast.cgi) searches for homologs in the rest of the OMM12 closed genome assemblies (Amur: GenBank accession CP065321, Amuc: CP065322, Bcae: CP065319, Bani: CP065311, Bpse: CP065312, Cinn: CP065320, Eclo: CP065314, Efae: CP065317, Fpla: CP065315, Lreu: CP065318, Mint: CP065316, Tmur: CP065313). Protein alignments were performed using MAFFT^86^ and reordered based on similarity to the top protein in the alignment. Alignments of the peptides were visualized using the R package ggmsa^87^ (RRID:SCR_025175).

#### Tetramer generation and testing

Peptides were synthesized at GenScript to a purity of 85% and sent to the NIH Tetramer Core Facility (supported by NIH contract 75N93020D00005) at Emory University for loading onto I-Ab MHC tetramers conjugated to PE or APC. Tetramers were initially tested for their capacity to bind to the cognate TCR on the reporter cell lines expressing them while simultaneously optimizing for temperature (4°C or 37°C) and length of staining (30 min or 1 hour) by staining first with the APC and PE conjugated tetramers together with CD16/32 Fc Block (Fc Block; clone 2.4G2, UCSF core, 1:200) in FACS buffer without azide containing dasatinib (final concentration 50 nM). Samples were stained with Aqua (1:1000), BUV395 anti-mouse CD45 (clone 30-F11, BD cat no. 564279, 1:400), BUV737 anti-mouse CD3 (clone 17A2, BD cat no. 612803, 1:400), BV785 anti-mouse CD4 (clone GK1.5, BioLegend cat no. 100453, 1:400), PerCP-Cy5.5 anti-mouse CD8 (clone 53-6.7, eBioscience cat no. 45-0081-82, 1:400), APC-eFluor780 anti-human CD4 (clone RPA-T4, eBioscience cat no. 47-0049-41, 1:400), and PE-Cy7 anti-mouse TCRβ (clone H57-597, BioLegend cat no. 109222, 1:200).

For T cell phenotyping assays, tissues were processed to single cell suspensions as described previously. Staining conditions chosen from optimization were used for staining with tetramer. Cells were incubated with paired APC and PE conjugated I-Ab-restricted tetramers, together with Fc Block and BV650 anti-mouse CXCR5 (clone L138D7, BioLegend cat no. 145517, 1:50) in FACS buffer without azide containing dasatinib for 1 hour at 37°C to preserve TCR surface expression during tetramer binding. Cells were then stained with NIR followed by surface staining with PE-Cy5 anti-mouse Ly6G (clone 1A8, BioLegend cat no. 127672), PE-Cy5 anti-mouse B220 (clone RA3-6B2, BioLegend cat no. 103210, 1:250), BV510 anti-mouse CD45 (clone 30-F11, BioLegend cat no. 103138, 1:400), BUV737 anti-mouse CD3ε (clone 145-2C11, BD cat no. 612771, 1:400), BV711 anti-mouse CD44 (clone IM7, BioLegend cat no. 103057, 1:400), BV785 anti-mouse CD4 (clone GK1.5, BioLegend cat no. 100453, 1:400), PerCP-Cy5.5 anti-mouse CD8α (clone 53-6.7, eBioscience cat no. 45-0081-82, 1:400), and BUV395 anti-mouse CD62L (clone MEL-14, BD cat no. 740218, 1:600). Cells were then fixed and permeabilized using the eBioscience Foxp3/Transcription Factor Staining Buffer Set (Thermo Fisher cat no. 00-5523-00) and stained intracellularly with BUV615 anti-mouse Bcl6 (clone K112-91, BD cat no. 568060, 1:100), FITC anti-mouse Foxp3 (clone FJK-16s, Invitrogen cat no. 53-5773-82, 1:500), PE-CF594 anti-mouse RORγt (clone Q31-378, BD cat no. 562684, 1:500), PE-Cy7 anti-mouse Tbet (clone 4B10, BioLegend cat no. 644824, 1:200), and eFluor 450 anti-mouse GATA3 (clone TWAJ, Invitrogen cat no. 48-9966-42, 1:100).

## Quantification and statistical analysis

All statistical tests for mouse T cell phenotyping experiments by flow cytometry were computed using GraphPad Prism and described in the Figure Legends. To characterize the phenotype of tetramer⁺ cells (Figures 4, 5, S4, and S5), we required a specific, above-background signal in each mouse. Only mice in which tetramer⁺ frequency exceeded the mean + 1 SD of the frequency observed in germ-free (GF) or Altered Schaedler Flora (ASF)-colonized control mice (as appropriate for each tetramer) were included in phenotype distribution analyses; mice that did not meet this threshold were excluded because tetramer⁺ events in these animals could not be reliably distinguished from background staining. Tetramer⁺ frequency itself is reported for all mice analyzed, regardless of this threshold.

## Use of artificial intelligence (AI)-assisted technologies

AI (Anthropic Claude) assisted in writing code used to extract data from Seurat objects for use in figures and to iterate through plasmid files for supplemental tables. All outputs were validated by the researchers.

## SUPPLEMENTAL FIGURES

**Figure S1.**
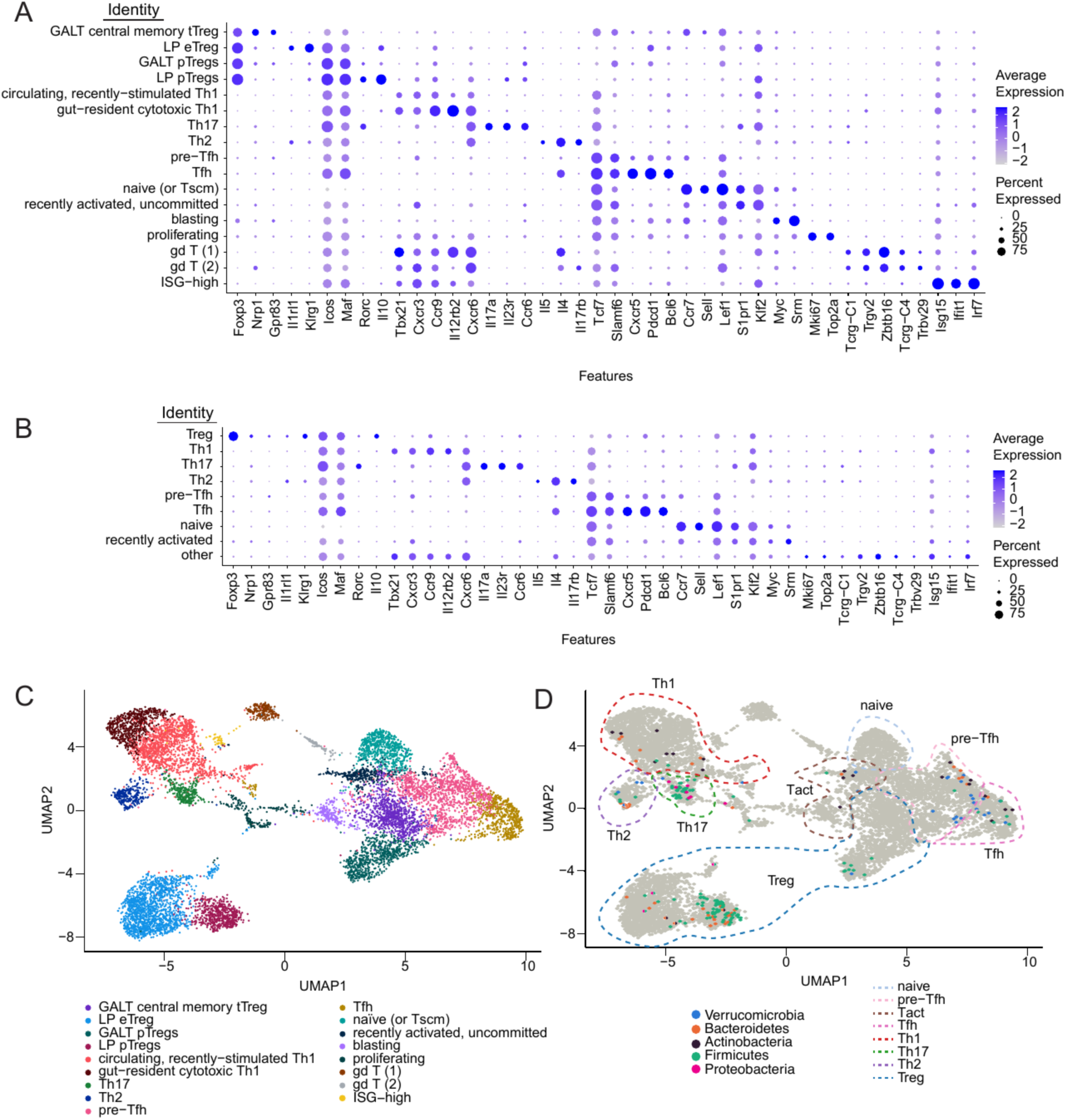
Marker-gene validation of T cell phenotype calls for single cell clusters, related to Figure 1. **(A)** Dot plot of relevant gene expression used to define Th clusters shown in Figure S1D. (B) Dot plot of relevant gene expression in manually-merged, broad Th clusters related to Figure 1D-F. (C) UMAP of scRNA-seq cluster assignments based on key marker expression shown in S1A and described in Methods. Tscm = T stem-like central memory, ISG=Interferon stimulated genes. (D) UMAP showing location of phyla level specificity of cells expressing microbiota-reactive TCRs. Broadly categorized T cell clusters shown in 1D are indicated with dashed lines and labeled.

**Figure S2.**
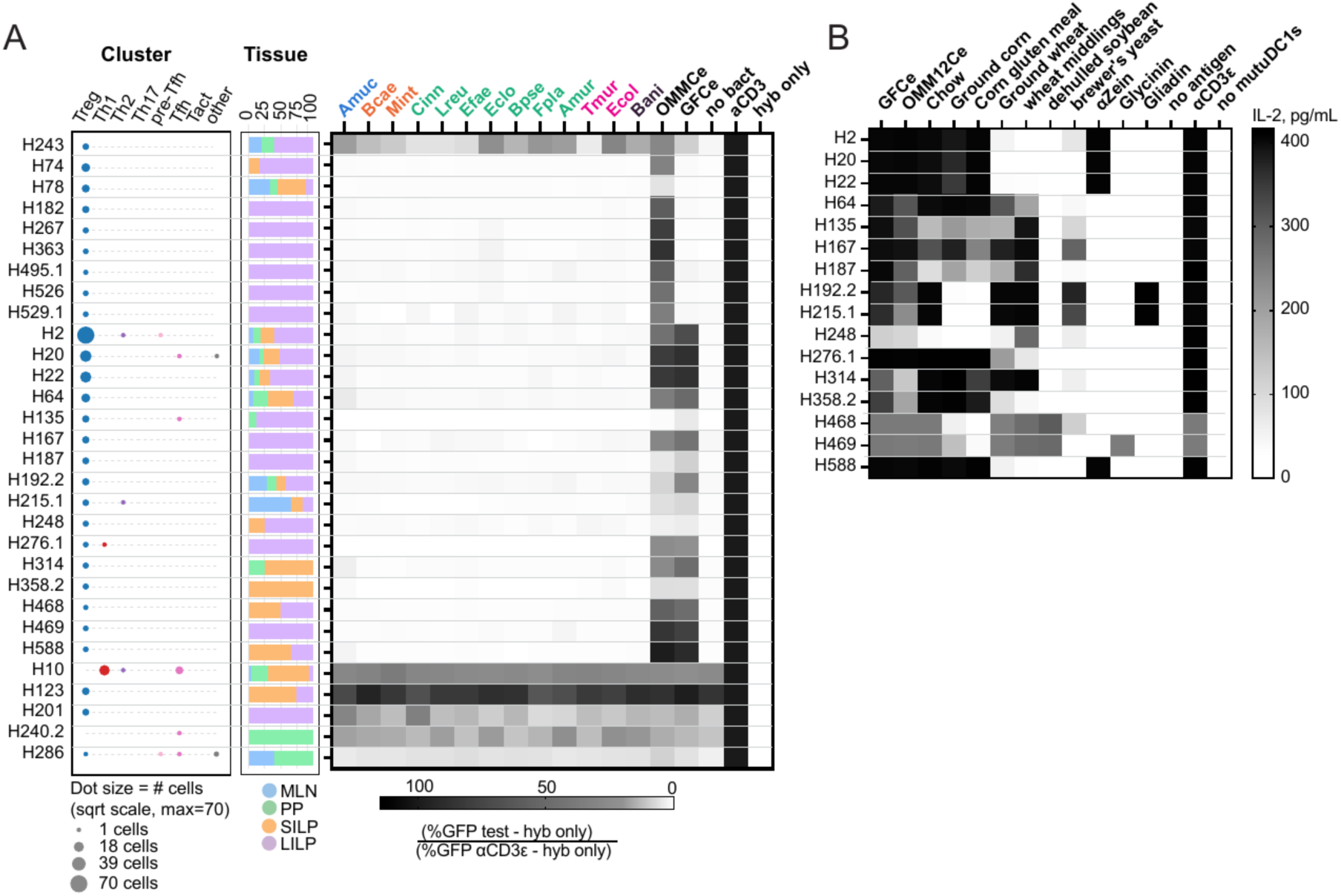
Additional microbiota/dietary TCR reactivity screening, related to Figure 1. **(A)** Compilation of Th phenotype (left), tissue origin (middle) and reactivity (right) data for T cells with reactivity that could not be mapped to individual OMM12 members (OMMCe+ = microbiota-reactive but not mappable to a single microbe, GFCe+ and OMMCe+ = likely food or self). (B) Specificity of food-reactive TCRs to components of mouse chow or known food peptides. αZein peptide (FYQQPIIGGAL), Glycinin peptide (EYVSFKTNDT), gliadin peptide (CNVYIPPYCTIAP).

**Figure S3.**
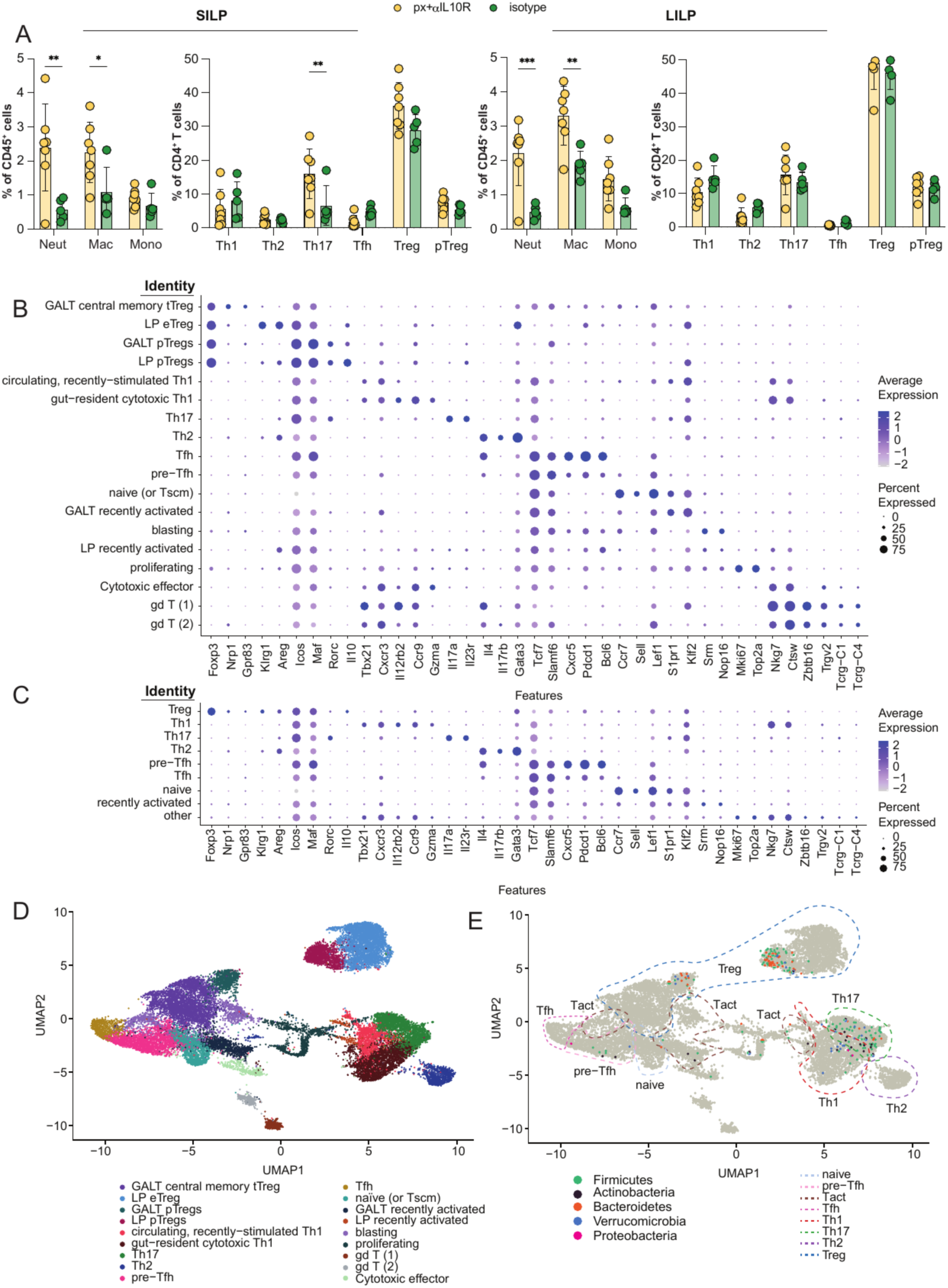
Validation of intestinal inflammation and T cell phenotype clusters by marker gene expression, related to Figure 2. (A) Summary of the frequencies of myeloid (neutrophils(Neut), macrophages(mac), and monocytes(mono)) as well as T helper(Th) cells collected at d17 from the SILP (Left) and LILP (Right) of the indicated groups of mice. Statistical significance was determined by two-way ANOVA with Sidak’s multiple comparisons test. *p < 0.05, **p < 0.01, ***p < 0.001. (B) Dot plot of relevant gene expression used to define Th clusters shown in Figure S3D. (C) Dot plot of relevant gene expression of manually-merged broad Th phenotype clusters related to Figure 2E. (D) UMAP of scRNA-seq cluster assignments based on key marker expression (described in Methods). (E) UMAP showing location of phyla level specificity of cells expressing microbiota-reactive TCRs. Broadly categorized T cell clusters shown in Figure 1B are indicated with dashed lines and labeled.

**Figure S4.**
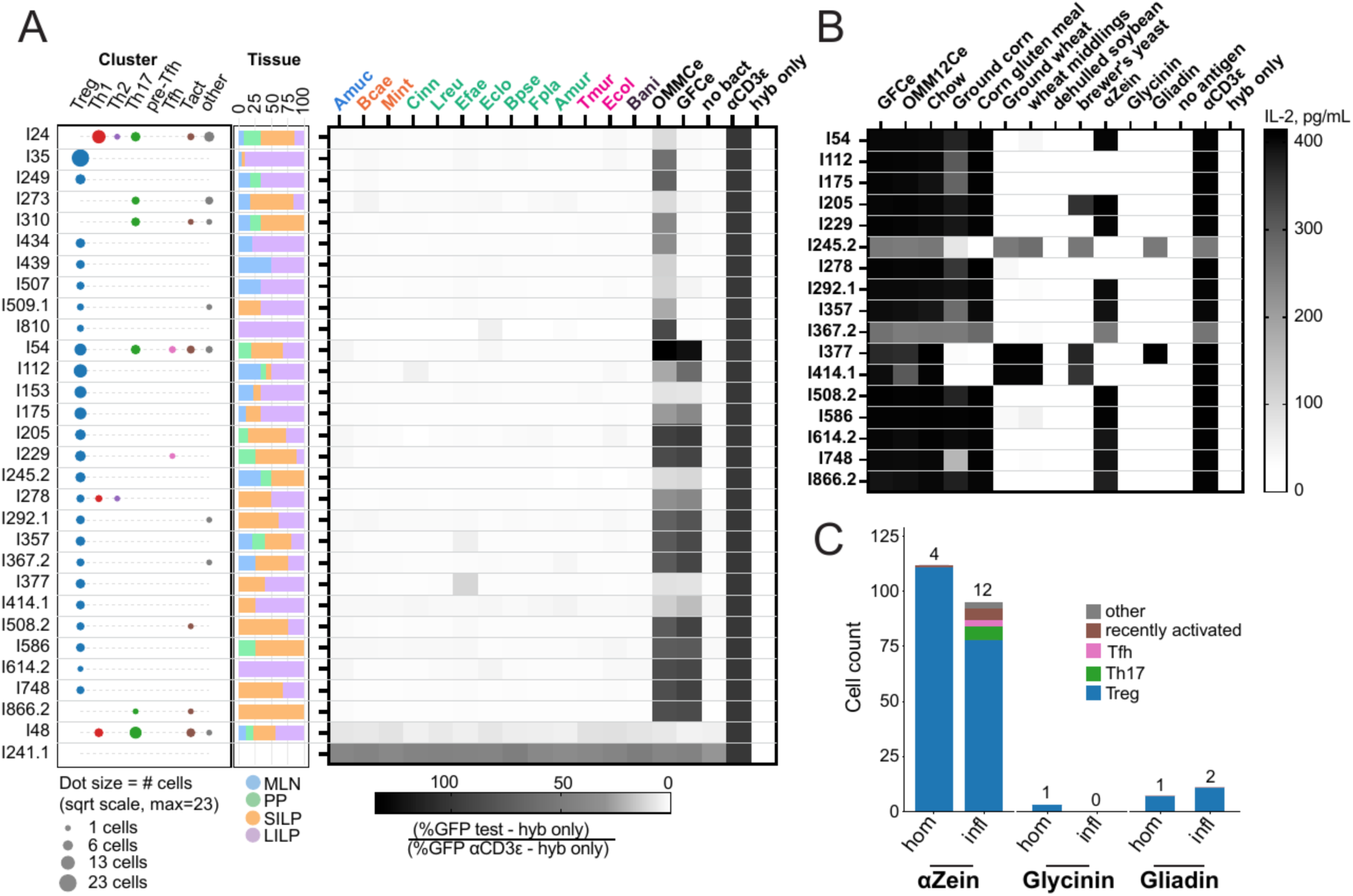
Additional microbiota/dietary TCR reactivity screening, related to Figure 2. (A) Compilation of Th phenotype (left), tissue origin (middle) and reactivity (right) data for T cells with reactivity that could not be mapped to individual OMM12 members (OMMCe+ = microbiota-reactive but not mappable to a single microbe, GFCe+ and OMMCe+ = probably food, or self). (B) Specificity of food-reactive TCRs to components of mouse chow or known food peptides. (C) Phenotypes of cells present in the homeostasis and inflammation single cell datasets that recognize αZein, glycinin and gliadin peptides. Note: inflammation clonotype 17 (I17) was identical to homeostasis clonotype 2 and therefore I17 is not included in S4A and S4B but is included in the cell counts for αZein-reactive cells during inflammation.

**Figure S5.**
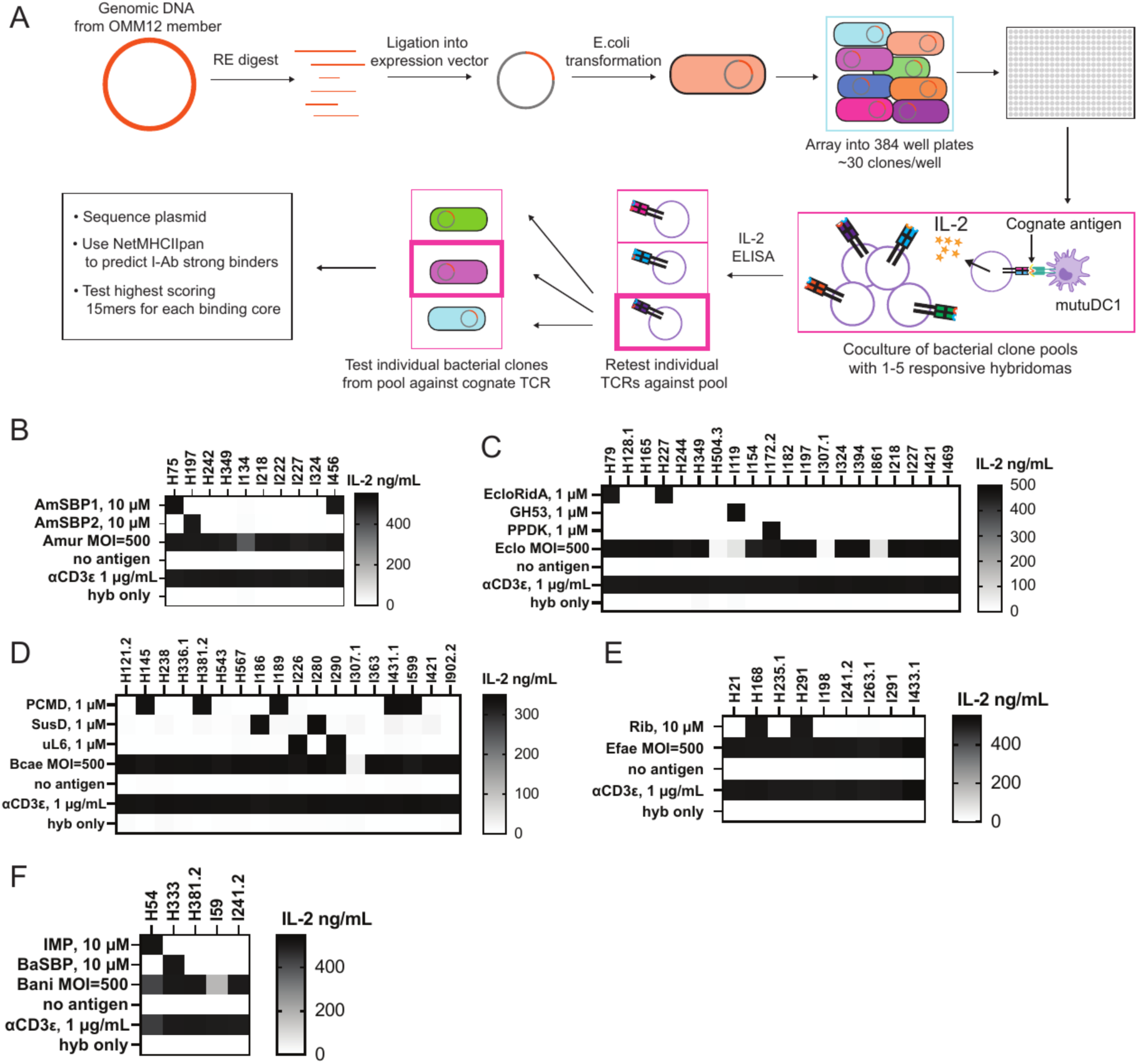
Peptide discovery pipeline and validation of peptide stimulation of additional TCRs, related to Figure 3. (A) Overview of antigen discovery pipeline. Genomic DNA from one of the OMM12 microbes was isolated and partially digested to generate a range of fragment sizes. The DNA fragments were ligated into an IPTG-inducible expression vector to create a genomic (g)DNA library. The gDNA library was transformed into E.coli and pools of ∼30 clones were arrayed in 2xYT+carb in 384 deep well plates. Overnight cultures were diluted, grown to log phase and then protein expression was induced overnight at room temperature (RT). Supernatants were removed and bacteria were transferred in PBS to 384 PCR plates to be heat-killed in a thermocycler. Pools of microbiota-reactive TCRs were cocultured with the heat-killed bacteria and mutuDC1s overnight. Supernatants from the coculture assay were assessed for the presence of IL-2 by ELISA. If a well was positive for IL-2, individual TCR hybridomas were screened for reactivity against the pooled bacteria. Then the pool-reactive TCRs were cocultured with individual heat-killed bacteria colonies from the pool. Plasmids were isolated from stimulatory bacteria and sent for whole plasmid sequencing. Any protein sequence present in the plasmid insert was searched for protein coding sequences that were scored for H2 Ab binding likelihood. Strong binding peptides were tested for their ability to stimulate in coculture with the TCR hybridoma and mutuDC1s. (B-F) Heat maps of reactivity measured by IL-2 production are shown for (B) Amuris-, (C) Eclo-, (D) Bcae-, (E) Efae- and (F) Bani-reactive TCRs in coculture assays.

**Figure S6.**
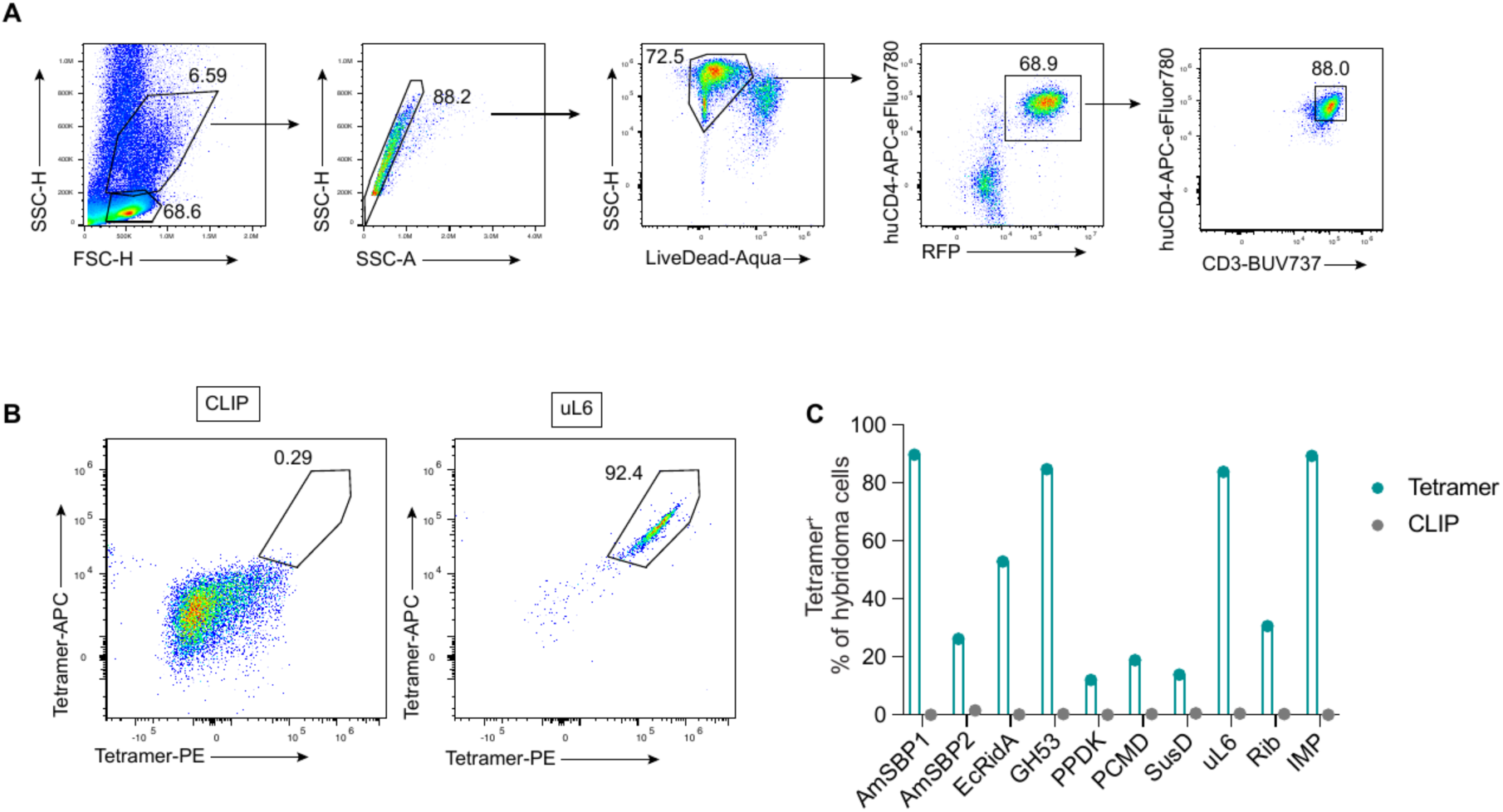
Validation of MHC class II tetramer specificity. (A) Gating strategy for identifying tetramer-stained hybridoma cells. Hybridomas were gated on FSC-H/SSC-H, singlets (SSC-H/SSC-A), viable cells (LiveDead-Aqua⁻), huCD4⁺RFP⁺, and CD3⁺huCD4⁺ prior to tetramer staining. (B) Representative dual-color tetramer staining (Tetramer-PE vs. Tetramer-APC) of the uL6-specific hybridoma, gated on the CD3⁺huCD4⁺ population in (A), stained with a CLIP-loaded negative-control tetramer (left) or the cognate uL6 tetramer (right).(C) Quantification of tetramer specificity across all ten epitope-specific hybridomas. Percentage of hybridoma cells stained by the cognate tetramer (teal) versus the CLIP-loaded negative-control tetramer (gray) for AmSBP1, AmSBP2, EcRidA, GH53, PPDK, PCMD, SusD, uL6, Rib, and IMP.

**Figure S7.**
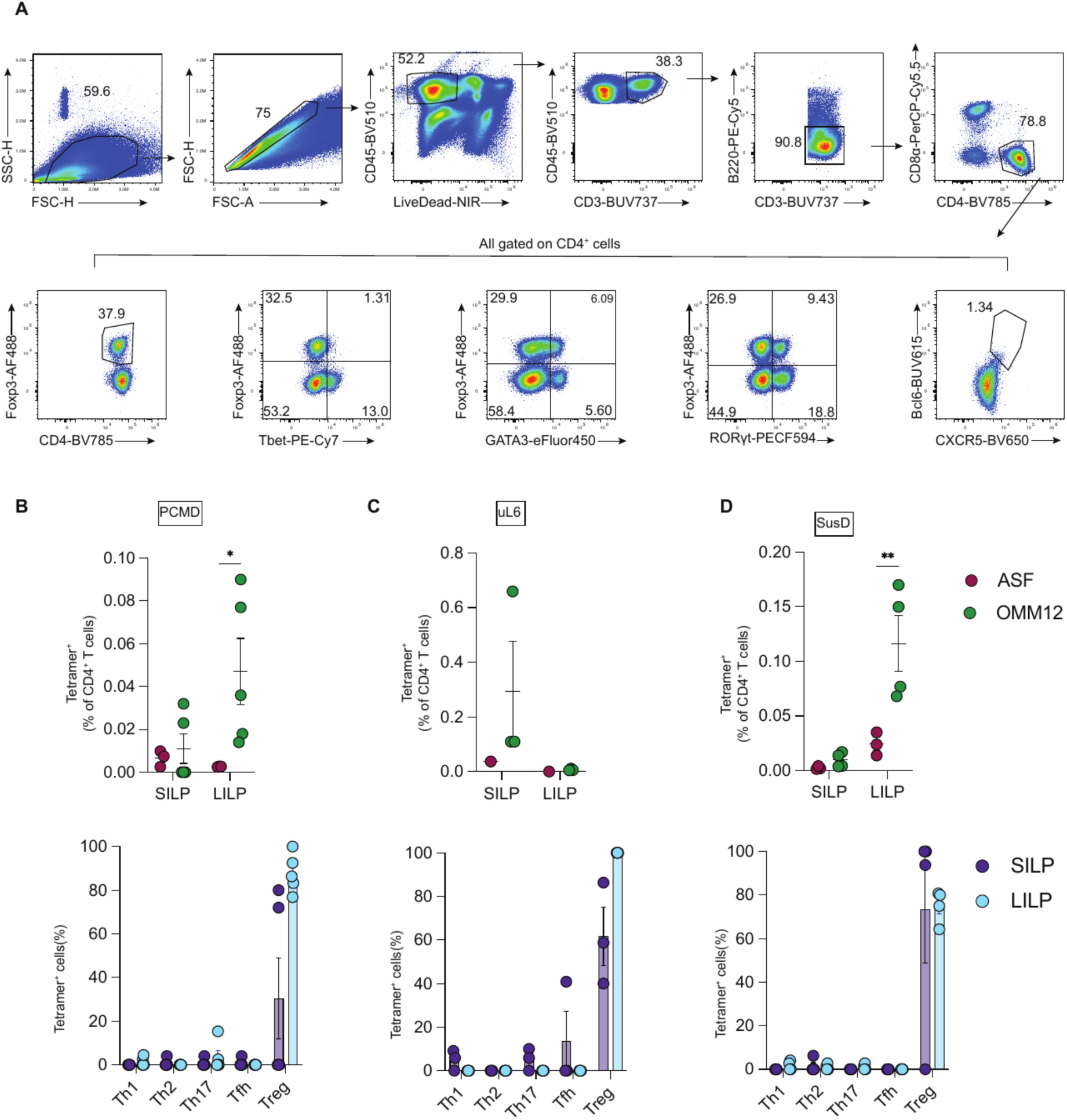
Gating strategy for identifying antigen-specific CD4⁺ T cells in vivo and validation of Bcae-tetramers individually. (A) Gating strategy for identifying tetramer⁺ CD4⁺ T cell subsets in vivo. Cells were gated on FSC-H/FSC-A, singlets, viable CD45⁺ cells, CD3⁺, B220⁻, and CD4⁺CD8α⁻ T cells. The tetramer⁺ gate and all subset-defining gates (from left to right): Foxp3⁺ (Treg), Tbet⁺Foxp3⁻ (Th1), GATA3⁺Foxp3⁻ (Th2), RORγt⁺Foxp3⁻ (Th17), and Bcl6⁺CXCR5⁺ (Tfh), were drawn on the CD4⁺ T cell population. The Th gates were subsequently mirrored onto the tetramer⁺ gate. (B–D) Frequency of individual Bcae-tetramer⁺ cells among CD4⁺ T cells in the small intestinal (SILP) and large intestinal (LILP) lamina propria of ASF and OMM12 mice (top), and phenotype distribution of tetramer⁺ cells in each tissue (bottom). Each dot represents an individual mouse. Statistical comparisons were not performed for (C) due to insufficient sample size in some groups. For (B,D) tetramer frequency statistical significance was determined by two-way ANOVA with Sidak’s multiple comparisons test; *p < 0.05, **p < 0.01.

**Figure S8.**
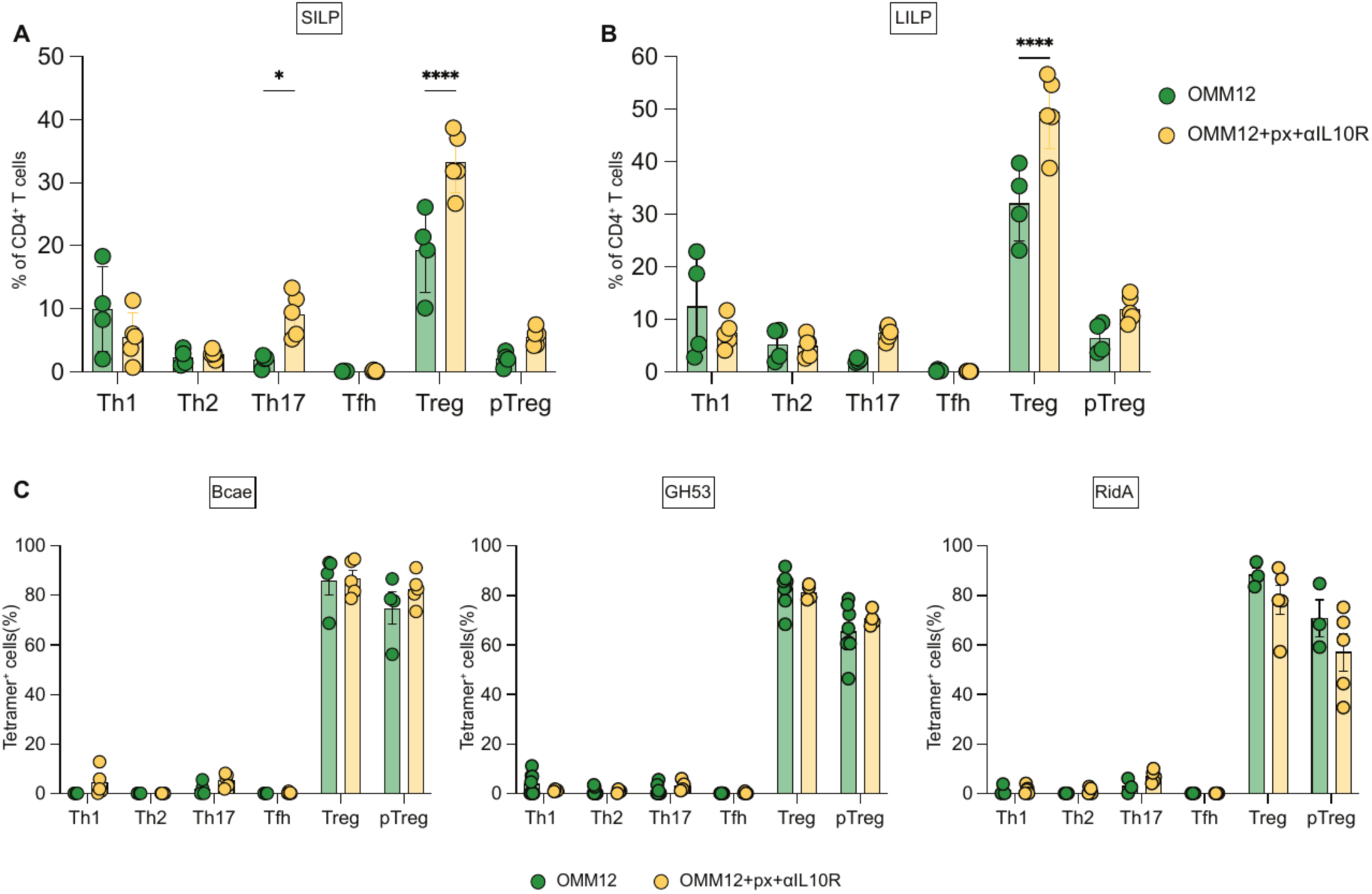
Characterization of Piroxicam and IL-10R blockade and additional tetramer phenotyping, related to Figure 4. (A, B) Phenotype distribution of endogenous CD4⁺ T cells in the SILP (A) and LILP (B) of OMM12 vs. OMM12+piroxicam+anti-IL-10R treated mice. (C) Phenotype distribution of tetramer⁺ CD4⁺ T cells from the LILP for the pooled Bcae tetramer, GH53, and RidA, comparing OMM12 and OMM12+piroxicam+anti-IL-10R treated mice. For (C) and (D), only mice with tetramer⁺ frequency exceeding the mean + 1 SD of the ASF control frequency were included (see Methods). Data are shown as mean ± SEM. Statistical significance was determined by two-way ANOVA with Sidak’s multiple comparisons test; *p < 0.05, ****p < 0.0001.

**Figure S9.**
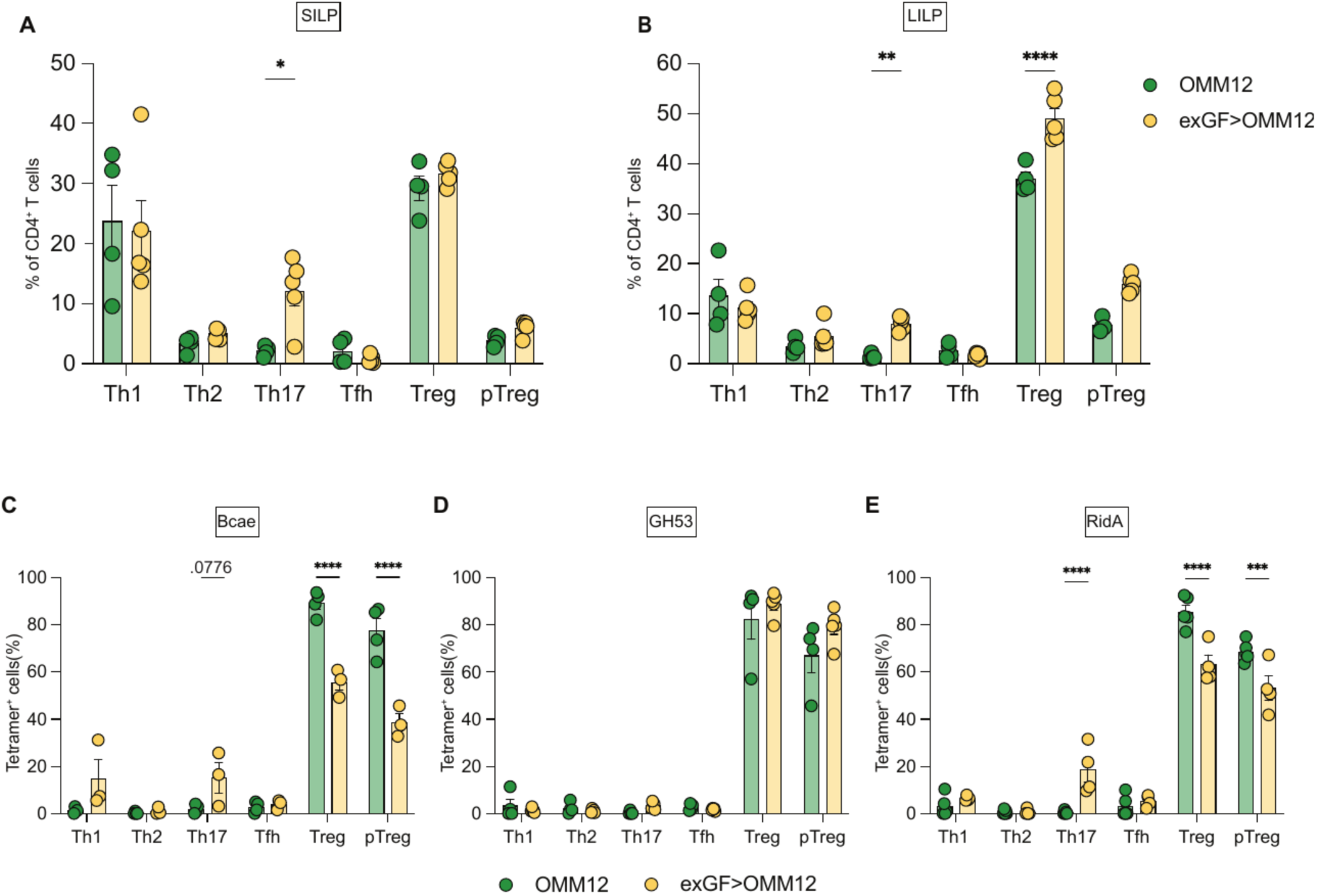
Horizontal colonization alters antigen-specific T cell phenotype in an epitope-dependent manner, related to Figure 5. (A, B) Phenotype distribution of endogenous CD4⁺ T cells in the SILP (A) and LILP (B) of OMM12 vs. exGF>OMM12 mice. (C–E) Phenotype distribution of tetramer⁺ CD4⁺ T cells from the LILP for the pooled Bcae tetramer (C), GH53 (D), and RidA (E), comparing OMM12 and exGF>OMM12 mice. Only mice with tetramer⁺ frequency exceeding the mean + 1 SD of the ASF (C, D) or GF (E) control frequency were included (see Methods). Data are shown as mean ± SEM. Statistical significance was determined by two-way ANOVA with Sidak’s multiple comparisons test; *p < 0.05, **p < 0.01, ***p < 0.001, ****p < 0.0001.

## SUPPLEMENTAL TABLES

**Table S1. | TRAV and TRBV plasmid sequences used in the generation of Sleeping Beauty Transposon-TCR constructs.** All mouse TRAV and TRBV sequences that were cloned into the Sleeping Beauty Transposon or pCR2-Blunt II-TOPO vector, respectively. Description of columns is as follows: Variable gene name: Cloned TCR alpha or beta variable gene name, Notes: any differences notes from 10x reference genome, Variable gene sequence = Variable gene sequence in plasmid, Note: Shared plasmid backbone sequences are listed as separate entries with [TRAV] or [TRBV] indicating where the additional TRAV or TRBV sequence is inserted.

**Table S2. | All cloned TCR sequences and reactivity metadata.** Sequences of all TCRs that were reactive in at least one condition and all information associated with that reactivity. Description of columns is as follows: TCR ID = name of plasmid. Condition = 10x dataset. Clonotype = Cellranger assigned clonotype. TRAV Consensus Used = Cellranger assigned consensus sequence name. TRBV Consensus Used = Cellranger assigned consensus sequence name. Sequence = Nucleotide sequence in the plasmid. TRAV = TRAV name in TCR. CDR3a = amino acid sequence of CDR3a. TRBV = TRBV name in TCR, CDR3b = amino acid sequence of CDR3b, Source = Species of bacteria or other source (i.e. Chow) that the TCR reacts with. (Mmus = self, NE = cloning attempted but not expressed, NR = tested and not reactive under any tested conditions, GFCe = Ceca contents from germ-free mice, OMMCe = Ceca contents from OMM12-colonized mice, underscore used between reactive sources). Protein = Protein name if identified (UNK = Unknown), Peptide = Peptide amino acid sequence if identified (UNK = unknown). gDNA screened = Included in any bacterial genomic DNA library screen. Confidence = Filter for inclusion in any plots of reactivity (LOW = requires followup, HIGH = confirmed in at least 2 coculture experiments, if underscore present then order corresponds to Source). TRAV = TRAV in consensus. TRAJ = TRAJ in consensus. CDR3a (aa) = amino acid sequence of CDR3a. CDR3a nt (contig) = nucleotide sequence of CDR3a. TRBV = TRBV in consensus. TRBJ = TRBJ in consensus. CDR3b (aa) = amino acid sequence of CDR3b. CDR3b nt (contig) = nucleotide sequence of CDR3b. TCRa nt (from construct) = TCRa sequence present in plasmid. TCRa matches contig = If TRUE, TCR construct and consensus match exactly, if FALSE, TCR construct differs from consensus. TCRb nt (from construct) = TCRb sequence present in plasmid. TCRb matches contig = If TRUE, TCR construct and consensus match exactly, if FALSE, TCR construct differs from consensus. Mismatch details = explanation of why a difference exists (e.g. used nearly identical TRAV/TRBV, nucleotide sequence changed to accommodate cloning).

## Notes

### Competing Interest Statement

The authors have declared no competing interest.

